# Comparison of whole-genome assemblies of European river lamprey (*Lampetra fluviatilis*) and brook lamprey (*Lampetra planeri*)

**DOI:** 10.1101/2024.12.06.627158

**Authors:** Ole K. Tørresen, Benedicte Garmann-Aarhus, Siv Nam Khang Hoff, Sissel Jentoft, Mikael Svensson, Eivind Schartum, Ave Tooming-Klunderud, Morten Skage, Anders Krabberød, Leif Asbjørn Vøllestad, Kjetill S. Jakobsen

## Abstract

We present haplotype-resolved whole-genome assemblies from one individual European river lamprey (*Lampetra fluviatilis*) and one individual brook lamprey (*Lampetra planeri*), usually regarded as sister species. The genome assembly of *L. fluviatilis* consists of pseudo-haplotype one, spanning 1073 Mb and pseudo-haplotype two, spanning 963 Mb. Likewise for the *L. planeri* specimen, the genome assembly spans 1049 Mb and 960 Mb for pseudo-haplotypes one and two, respectively. Both the *L. fluviatilis* pseudo-haplotypes have been scaffolded into 82 pseudo-chromosomes, with the same number for the *L. planeri* pseudo-haplotypes. All four pseudo-haplotype assemblies were annotated, identifying 21,479 and 16,973 genes in pseudo-haplotypes one and two for *L. fluviatilis*, and 24,961 and 21,668 genes in pseudo-haplotypes one and two for *L. planeri*. A comparison of the genomes of *L. fluviatilis* and *L. planeri*, alongside a separate chromosome level assembly of *L. fluviatilis* from the UK, indicates that they form a species complex, potentially representing distinct ecotypes. This is further supported by phylogenetic analyses of the three reference *Lampetra* genomes in addition to sea lamprey (*Petromyzon marinus*).

## Introduction

Freshwater fishes reside in lakes, rivers, and streams and often migrate between different habitats, such as within and between rivers and lakes. Diadromous fishes can also sometimes migrate between freshwater and marine environments. In particular, many species in postglacial lakes show large phenotypic plasticity and also possess many morphotypes – sometimes regarded as different species. Determining what constitutes a species has been challenging for many freshwater fishes; a typical example is Salmoniformes, such as trout, charr, and whitefish (Whiteley et al., 2019). The genetic structuring following glaciations and subsequent post-glacial invasions, together with phenotypic plasticity, has led to large among-population variation in morphology, behaviour and life history.

In Petromyzontidae lampreys, this has led to the evolution of so-called species pairs consisting of closely-related large migratory parasitic and non-parasitic freshwater-resident species (Docker 2009). The migratory and parasitic European river lamprey (*Lampetra fluviatilis)* and the non-migratory and non-parasitic brook lamprey (*Lampetra planeri*) are regarded as sister species. They have been the subject of several genetic studies, using mtDNA (mitochondrial DNA) (Bracken et al., 2015; Cahsan et al., 2020), RADseq (restriction-site associated DNA sequencing) (Hume et al., 2018; Mateus et al., 2013; Rougemont et al., 2017), and microsatellite markers (Rougemont et al., 2015). There is nonetheless no consensus if these two taxa are separate species, or merely ecotypes, with different life-history traits.

While *L. fluviatilis* and *L. planeri* are morphologically and behaviourally similar in their larval stages, sustaining themselves through filter feeding at the bottom of freshwater streams for the first five to seven years of their lives (Potter et al., 2015; Rougemont et al., 2015), they differ greatly upon entering maturity. When maturing, *L. planeri* develops eyes and the characteristic lamprey sucker mouth, degenerates its gut and stops feeding, only to then mate and die in the freshwater where it has spent its entire life (Rougemont et al., 2015). In contrast, following metamorphosis, *L. fluviatilis* enters a migratory and often anadromous, parasitic juvenile life stage, where it migrates to lakes or the sea to feed on larger fish. For up to three years, the juvenile *L. fluviatilis* lives as a parasite (Kelly and King, 2001; Rougemont et al., 2016) and returns at sexual maturity to running water to mate and die (Kelly and King, 2001; Rougemont et al., 2016). A central unanswered question is whether the morphological and life-history differences between the two species are due to genetics or phenotypic plasticity.

Genetic studies to date have not clearly identified any distinctions that would suggest two separate species or morphologically and behaviorally diverged ecotypes. It is thus suggested that the *L. fluviatilis/L. planeri* species pair is at different stages of speciation in different locations (Mateus et al., 2016; Rougemont et al., 2017). Therefore, whole genome sequencing at the population level needs to be performed to capture not only SNP (single nucleotide polymorphism) variation but also structural variation, such as chromosomal rearrangements, inversions, CNVs (copy-number variations) and STR (short tandem repeat) length variations. These investigations require high-quality reference genomes for the two sister species.

Here, we report two pseudo-haplotype resolved, chromosome-level reference genomes of *L. fluviatilis* and *L. planeri* (the first for this species), using long-read PacBio HiFi sequencing and scaffolding with Hi-C to achieve the standards of the Earth BioGenome Project (Lewin et al., 2022). The differences between the genome assemblies for the two species and two published chromosome-level assemblies of *L. fluviatilis* and *Petromyzon marinus* were investigated by phylogenetic and chromosomal synteny analyses and showed that the sister species were highly similar - likely forming a species complex. The new reference genomes will be ideal for future larger population genomic analyses to fully resolve the species versus ecotype question.

## Methods

### Sample acquisition and DNA extraction

In this study, two lamprey specimens – an *L. fluviatilis* and an *L. planeri* – were collected from different locations in Scandinavia. The *L. fluviatilis* specimen was caught in Åsdalsåa, Telemark, Norway (59.410917, 9.305889) on 2021.04.21 using electrofishing and transported live to the University of Oslo. The individual was euthanized in the laboratory using an overdose of methanesulfonate (MS-222) and decapitation. The fish was 170 mm long, and muscle, blood and heart tissues were extracted and snap-frozen in individual Eppendorf tubes using liquid nitrogen. Similarly, the *L. planeri* specimen was caught in Hunserödsbäcken, Skåne, Sweden (56.250944, 13.001400) on 2020.10.27 using electrofishing and euthanized on-site. The whole body was stored in 96% ethanol and subsequently shipped to Oslo. The fish was 122 mm long, and muscle, skin tissue, gill filaments, and the entire heart were dissected. All tissues from both lampreys were transferred to the Norwegian Sequencing Centre for library preparation and stored at −80 degrees C.

### Library preparation and sequencing for *de-novo* assembly

For PacBio HiFi sequencing, DNA was isolated from the *L. fluviatilis*’s blood and from the *L. planeri*’s muscle and skin tissue. For the *L. fluviatilis*, 10-20 µl of fresh blood was used per reaction, and the Circulomics Nanobind CBB Big DNA kit was applied with the blood and tissue protocol, following manufacturer guidelines. The high molecular weight DNA was eluted from the Nanodisk with 150µl Tris-Cl buffer and incubated overnight at room temperature. The resulting DNA was then quality-checked for its amount, purity, and integrity using UV-absorbance ratios, a Qubit BR DNA quantification assay kit, and a Fragment Analyzer with a DNA HS 50 kb large fragment kit. In contrast, for the *L. planeri*, 30 mg of dry-blotted, EtOH-stored muscle and skin tissue was used per reaction. The *L. planeri* followed the same isolation process as the *L. fluviatilis* with some additional steps: incubation with proteinase K for two hours at room temperature, followed by incubation with RNAse for an additional 30 minutes at the same temperature. The same quality assessment methods were then applied to the isolated DNA of *L. fluviatilis*.

DNA from both the *L. fluviatilis* and *L. planeri* underwent PacBio HiFi sequencing by the Norwegian Sequencing Centre using three 8M SMRT cells on PacBio Sequel II after a size selection using the BluePippin system with an 11 kb cut-off (Wang et al., 2021). For the *L. fluviatilis*, two libraries were created from muscle tissue; while for the *L. planeri*, two libraries were prepared from muscle and skin tissues.

Both the *L. fluviatilis* and *L. planeri* samples underwent Hi-C sequencing to capture their three-dimensional chromatin structures. For the *L. fluviatilis* specimen, the library preparation followed the “Omni-C Proximity Ligation assay for Non-mammalian samples, version 1.0” protocol from the manufacturer. This involved grinding 20 mg of fresh, snap-frozen heart tissue to a fine powder, followed by lysis and proximity ligation. The prepared library was then sequenced on a NovaSeq 6000 Sequencing System at the Norwegian Sequencing Centre, using one full S Prime NovaSeq Flow Cell for 2 x 150 bp paired-end sequencing.

Similarly, for the *L. planeri*, 100 mg of gill tissue stored in ethanol was used. The library was prepared using an “Arima Genome-Wide HiC+ Kit” and the “Arima-HiC 2.0 kit standard user guide for Animal tissues”-protocol. The sequencing was carried out on a NovaSeq 6000 at the Norwegian Sequencing Centre, utilizing one quarter of a NovaSeq Flow Cell for 2 x 150 bp paired-end sequencing.

### Genome assembly and curation

A full list of relevant software tools and versions is presented in Supplementary Table 1. We assembled the species using a pre-release of the EBP-Nor genome assembly pipeline (https://github.com/ebp-nor/GenomeAssembly). *KMC* (Kokot et al., 2017) was used to count k-mers of size 21 in the PacBio HiFi reads, excluding k-mers occurring more than 10,000 times. *GenomeScope* (Ranallo-Benavidez et al., 2020) was run on the k-mer histogram output from *KMC* to estimate genome size, heterozygosity, and repetitiveness. Ploidy level was investigated using *Smudgeplot* (Ranallo-Benavidez et al., 2020).

*HiFiAdapterFilt* (Sim et al., 2022) was applied on the HiFi reads to remove possible remnant PacBio adapter sequences. The filtered HiFi reads were assembled using *hifiasm* (Cheng et al., 2021) with Hi-C integration resulting in a pair of haplotype-resolved assemblies, pseudo-haplotype one (hap1) and pseudo-haplotype two (hap2) for each species. Unique k-mers in each assembly/pseudo-haplotype were identified using *meryl* (Rhie et al., 2020) and used to create two sets of Hi-C reads, one without any k-mers occurring uniquely in hap1 and the other without k-mers occurring uniquely in hap2. These k-mer filtered Hi-C reads were then aligned to each assembly using *BWA-MEM* (Li, 2013) with `-5SPM` options. If there are large scale structural differences between the pseudo-haplotypes, such as inversions, using the whole Hi-C dataset could enforce the wrong orientation in an inversion for instance. Filtering the dataset aims to avoid enforcing the wrong topology on the chromosomes.

The alignments were sorted based on name using samtools (Li et al., 2009) before applying *samtools fixmate* to remove unmapped reads and secondary alignments and to add a mate score, along with *samtools markdup* to remove duplicates. The resulting BAM files were used to scaffold the two assemblies using *YaHS* (Zhou et al., 2022) with the default options. *FCS-GX* (Astashyn et al., 2023) was used to search for contamination in the scaffolds. Contaminated sequences were removed. If a contaminant was detected at the start or end of a sequence, the sequence was trimmed using a combination of *samtools faidx, bedtools* (Quinlan and Hall, 2010) *complement*, and *bedtools getfasta*. If the contaminant was internal, it was masked using *bedtools maskfasta*. The mitochondrion was searched for in contigs and reads using *MitoHiFi* (Uliano-Silva et al., 2023).

The assemblies were manually curated using *PretextView*, merging sequences that were supported by Hi-C signals and breaking some where the signal was lacking. Chromosomes were identified by inspecting the Hi-C contact map in *PretextView* and named according to homology to kcLamFluv1.

### Genome annotation

We annotated the genome assemblies using a pre-release version of the EBP-Nor genome annotation pipeline (https://github.com/ebp-nor/GenomeAnnotation). In general, default options were used for the different tools, but the specific parameters are detailed in the pipeline. First, *AGAT* (https://zenodo.org/record/7255559) *agat_sp_keep_longest_isoform*.*pl* and *agat_sp_extract_sequences*.*pl* was used on the *P. marinus* (GCA_010993605.1) genome assembly and annotation to generate one protein (the longest isoform) per gene. *Miniprot* (Li, 2023) was used to align the proteins to the curated assemblies. UniProtKB/Swiss-Prot (Consortium et al., 2022) release 2022_03 and the Vertebrata part of OrthoDB v11 (Kuznetsov et al., 2022) were also aligned separately to the assemblies. *Red* (Girgis, 2015) was run via *redmask* (https://github.com/nextgenusfs/redmask) on the assemblies to mask repetitive areas. *GALBA* (Brůna et al., 2023; Buchfink et al., 2015; Hoff and Stanke, 2018; Li, 2023; Stanke et al., 2006) was run with the *P. marinus* proteins using the miniprot mode on the masked assemblies. The *funannotate-runEVM*.*py* script from *Funannotate* was used to run *EvidenceModeler* (Haas et al., 2008) on the alignments of *P. marinus* proteins, UniProtKB/Swiss-Prot proteins, Vertebrata proteins and the predicted genes from *GALBA*.

The resulting predicted proteins were compared to the protein repeats that *Funannotate* distributes using *DIAMOND blastp*; the predicted genes were filtered based on this comparison using *AGAT*. The resultant filtered proteins were compared to the UniProtKB/Swiss-Prot release 2022_03 using *DIAMOND* (Buchfink et al., 2015) *blastp* to find gene names, and *InterProScan* (Jones et al., 2014) was used to discover functional domains. *AGATs agat_sp_manage_functional_annotation*.*pl* was used to attach the gene names and functional annotations to the predicted genes. *EMBLmyGFF3* (Norling et al., 2018) was used to combine the fasta files and GFF3 files into an EMBL format for submission to ENA. We also annotated the *P. marinus* (kPetMar1; GCA_010993605.1) and another river lamprey (kcLamFluv1; GCA_964198585.1) using the same approach as described here.

### Evaluation of the assemblies and comparative genomics

All the evaluation tools have also been implemented in a pipeline, similar to assembly and annotation (https://github.com/ebp-nor/GenomeEvaluation). To evaluate the diploid assembly, we ran *Flagger* (Liao et al., 2023) to detect possible mis-assemblies. The HiFi reads were mapped to the diploid assembly (created by concatenating the two pseudo-haplotypes) using *winnowmap* (Jain et al., 2022). *Secphase* (Liao et al., 2023) was run on the BAM file produced by *winnowmap* to correct the alignments of the reads by scoring them based on marker consistency and selecting the alignment with the highest score as primary. SNPs were called from the corrected BAM file by *DeepVariant* (Poplin et al., 2018) using default parameters for PacBio HiFi data and filtered to keep only biallelic SNPs. *Flagger* (Liao et al., 2023) was then run on the corrected BAM file together with the filtered VCF and categorized the diploid assembly into erroneous, duplicated, haploid, collapsed, and unknown regions.

*Merqury* (Rhie et al., 2020) was used to assess the completeness and quality of the genome assemblies by comparing them to the k-mer content of both the Hi-C reads and PacBio HiFi reads. *BUSCO* (Manni et al., 2021) was used to assess the completeness of the genome assemblies by comparing against the expected gene content in the metazoa lineage. We also ran *BUSCO* on *P. marinus* (kPetMar1; GCA_010993605.1) and another river lamprey (kcLamFluv1; *L. fluviatilis;* GCA_964198585.1). *Gfastats* (Formenti et al., 2022) was used to output different statistics of the assemblies, including kPetMar1 and kcLamFluv1.

*BlobToolKit* and *BlobTools2* (Laetsch and Blaxter, 2017), in addition to *blobtk* were used to visualize assembly statistics. To generate the Hi-C contact map image, the Hi-C reads were mapped to the assemblies using *BWA-MEM* (Li, 2013) using the same approach as above. Finally, *PretextMap* was used to create a contact map which was visualized using *PretextSnapshot*.

To characterize the genomic differences between the different assemblies (both pseudo-haplotypes of both species, in addition to kcLamFluv1), we ran *nucmer* from the *MUMmer* (Marçais et al., 2018) genome alignment system on the homologous chromosomes from the assemblies, using these parameters `--maxmatch -l 100 -c 500`. The resulting alignments were processed with *dnadiff*, also from *MUMmer*, which produced reports listing the number of insertions, SNPs, and indels between the different assemblies. *EMBOSS* (Rice et al., 2000) *infoseq* was used to calculate GC content of the different sequences.

We ran *OrthoFinder* (Buchfink et al., 2015; Emms and Kelly, 2019, 2018, 2017) on the predicted proteins for all the assemblies to infer multiple sequence alignment gene trees. *OrthoFinder* was run with the option msa using *MAFFT* (Katoh and Standley, 2013) as the multiple alignment tool and *IQ-TREE* (Minh et al., 2020) for gene tree inference. We obtained the species tree from the gene trees using *ASTRAL-Pro3* (Zhang and Mirarab, 2022) by optimizing the objective function of *ASTRAL-Pro* (Zhang et al., 2020).

To inspect the syntenic relationship among the genomes between the different species, we ran *MCScanX* (Wang et al., 2012) and visualized the results using *Synvisio* (Bandi and Gutwin, 2020). First, we used *DIAMOND blastp* (v2.1.16) with the options `-q ${IN_PROT} -p 16 -e 1e-10 --max-hsps 5` with annotated proteins for kcLamPlan1.2.hap1, kcLamPlan1.2.hap2, kcLamFluv2.2.hap1, kcLamFluv2.2.hap2, kcLamFluv1, and kPetMar1 as input data. Subsequently, *MCScanX* was run with default settings, and results visualized using the online interactive platform *Synvisio*.

## Results

### De novo genome assembly and annotation

The genome from the European river lamprey (*L. fluviatilis*) had an estimated genome size of 742 Mb, with 1.09% heterozygosity and a bimodal distribution based on the k-mer spectrum (Supplementary Figure 1). The genome from the European brook lamprey (*L. planeri*) had an estimated size of 720 Mb, with 1.1% heterozygosity and a bimodal distribution based on its k-mer spectrum (Supplementary Figure 2.). A total of 38-fold coverage by the PacBio single-molecule HiFi long reads and 100-fold coverage by the Omni-C reads resulted in two haplotype-separated assemblies for *L. fluviatilis. L. planeri* was assembled with 44-fold PacBio and 120-fold Arima Hi-C reads.

For *L. fluviatilis*, the final assemblies had total lengths of 1073 Mb (Figure 1 and Table 1) and 963 Mb (Table 1 and Supplementary Figure 3) for pseudo-haplotypes one and two, respectively. For *L. planeri*, the pseudo-haplotypes one and two had total lengths of 1049 Mb (Figure 1 and Table 1) and 960 Mb (Table 1 and Supplementary Figure 3), respectively. Pseudo-haplotypes one and two for *L. fluviatilis* have scaffold N50 sizes of 13.1 Mb and 13.4 Mb, respectively, and contig N50 of 2.7 Mb and 2.9 Mb, respectively (Table 1). *L. planeri* have scaffold N50 size of 12.9 Mb in both pseudo-haplotype one and two, and contig N50 sizes of 2.8 Mb and 3.0 Mb, respectively. 82 automosomes were identified in both pseudo-haplotypes for *L. fluviatilis* (chromosomes named after kcLamFluv1) and 82 in both pseudo-haplotypes in *L. planeri* (chromosomes also named after kcLamFluv1).

**Table 1:** Genome data for *L. fluviatilis*, kcLamFluv2 and *L. planeri*, kcLamPlan1, including accession numbers and genome assembly and annotation metrics for both haplotypes for both species.

| Project accession data |  |  |  |  |
| --- | --- | --- | --- | --- |
| Species | *L. fluviatilis* |  | *L. planeri* |  |
| Specimen | kcLamFluv2 |  | kcLamPlan1 |  |
| NCBI taxonomy ID | 7748 |  | 7750 |  |
| BioProject | PRJEB77187 |  | PRJEB77192 |  |
| BioSample ID | SAMEA115797768 |  | SAMEA115802553 |  |
| Isolate information | Male, fin |  | Sex not provided, fin |  |
| Raw data accessions |  |  |  |  |
| PacBio HiFi reads | ERX12712303, ERX12712308, ERX12712309 | 3 PACBIO_SMRT (Sequel II) runs: 2.5 M reads, 38.5 Gbp | ERX12713797, ERX12713780, ERX12713807 | 3 PACBIO_SMRT (Sequel II) runs: 3.2 M reads, 44.0 Gbp |
| Hi-C Illumina reads | ERX12712501 | 1 ILLUMINA (Illumina NovaSeq S4) run: 334 M pairs of reads, 100.8 Gbp | ERX12714064 | 1 ILLUMINA (Illumina NovaSeq S4) run: 407 M pairs of reads, 122.9 Gbp |
| Genome assembly metrics |  |  |  |  |
| HiFi read coverage | 38 |  | 44 |  |
| Assembly accession | ERZ24889083 | ERZ24889084 | ERZ24889000 | ERZ24889001 |
| Assembly identifier | kcLamFluv2.2.hap1 | kcLamFluv2.2.hap2 | kcLamPlan1.2.hap1 | kcLamPlan1.2.hap2 |
| Span (Mb) | 1073 | 963 | 1049 | 960 |
| Number of chromosomes | 82 | 82 | 82 | 82 |
| Number of contigs | 3060 | 1828 | 3142 | 3066 |
| Contig N50 length (Mb) | 2.7 | 2.9 | 2.8 | 3.0 |
| Longest contig (Mb) | 22.2 | 21.8 | 26.2 | 16.7 |
| Number of gaps | 1327 | 942 | 1113 | 873 |
| Number of scaffolds | 1733 | 886 | 2029 | 2193 |
| Scaffold N50 length (Mb) | 13.1 | 13.4 | 12.9 | 12.9 |
| Longest scaffold (Mb) | 41.1 | 41.3 | 41.1 | 41.0 |
| Consensus quality (QV) compared to Hi-C (compared to HiFi) | 38.6766 (54.8795) | 40.4312 (56.0416) | 34.9015 (52.7149) | 28.9161 (52.4613) |
| Both assemblies | 39.4197 (55.3907) |  | 31.0849 (52.5919) |  |
| *k*-mer completeness (percentage; compared to HiFi) | 83.8059 (89.3051) | 80.3495 (86.2251) | 89.6766 (91.5393) | 84.8223 (87.4288) |
| Both assemblies | 92.194 (98.4232) |  | 96.4739 (98.2448) |  |
| *BUSCO* * | C:91.4%[S:88.5%, D:2.9%],F:4.6%, M:4.0%,n:954 | C:83.9%[S:82.4%, D:1.5%],F:3.5%,M:12.6%,n:954 | C:89.8%[S:86.4%, D:3.4%],F:4.4%,M:5.8%,n:954 | C:83.5%[S:77.5%, D:6.0%],F:3.1%,M:13.4%,n:954 |
| Percentage of assembly mapped to chromosomes | 90.44 | 95.90 | 91.43 | 91.21 |
| *Flagger* ** | H: 79.33%, D: 20.35%, E:0.0%, C:0.03% | H: 82.20%, D: 17.54%, E:0.0%, C:0.03% | H: 75.27%, D: 21.22%, E:3.2%, C:0.03% | H: 75.57%, D: 18.69%, E:5.5%, C:0.02% |
| Organelles<br>(identified in<br>the genome<br>assembly) | MT |  | MT |  |
| **Genome annotation metrics** |  |  |  |  |
| Number of<br>protein-coding<br>genes | 21,479 | 16,973 | 24,691 | 21,668 |
| Number of<br>protein-coding<br>genes with<br>functional<br>domain*** | 20,126 | 7875 | 12,006 | 20,026 |
| Number of<br>protein-coding<br>genes with gene<br>names**** | 13,217 | 11,576 | 13,227 | 13,244 |
| BUSCO* | C:89.0%[S:84.6%,<br>D:4.4%],F:3.2%,<br>M:7.8%,n:954 | C:82.4%[S:79.8%,D<br>:2.6%],F:2.5%,M:15<br>.1%,n:954 | C:89.2%[S:85.5%,<br>D:3.7%],F:3.1%,M:<br>7.7%,n:954 | C:82.6%[S:76.8%,<br>D:5.8%],F:2.5%,M<br>:14.9%,n:954 |

\* BUSCO scores are based on the metazoa BUSCO set using v5.4.7. C = complete [S = single copy, D = duplicated], F = fragmented, M = missing, n = number of orthologues in comparison.
\*\*Flagger scores H = haploid, D = duplicated, E = error, C = collapsed
\*\*\*Number of genes annotated with a functional domain as found by InterProScan
\*\*\*\*Number of genes that had a match against a named protein in UniProtKB/Swiss-Prot

**Figure 1:**
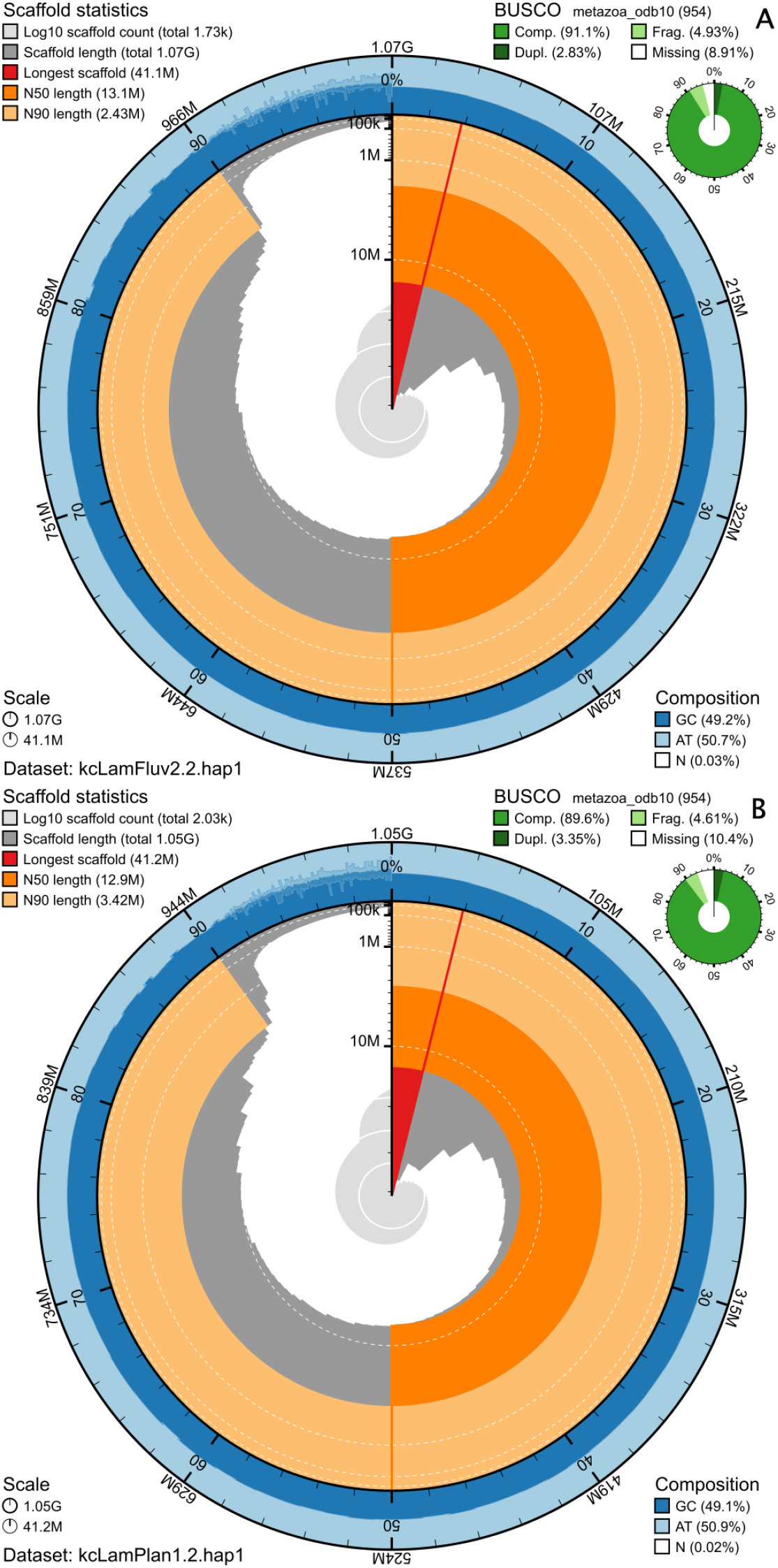
Metrics for the genome assemblies of *L. fluviatilis* (A) and *L. planeri* (B), pseudo-haplotype one for both species. The BlobToolKit Snailplots show N50 metrics and BUSCO gene completeness. The two outermost bands of the circle signify GC versus AT composition at 0.1% intervals, with mean, maximum and minimum. Light orange shows the N90 scaffold length, while the deeper orange is N50 scaffold length. The red line shows the size of the largest scaffold. All the scaffolds are arranged in a clockwise manner from largest to smallest, and shown in darker gray with white lines at different orders of magnitude, shown as a scale on the radius. The light gray shows the cumulative scaffold count. The scale inset in the lower left corner shows the total amount of sequence in the whole circle, and the fraction of the circle encompassed in the largest scaffold.

For *L. fluviatilis*, pseudo-haplotype one had 91.4%, and pseudo-haplotype two had 83.9% complete BUSCO genes using the metazoa lineage set. *L. planeri* pseudo-haplotype one had 89.8% and pseudo-haplotype two 83.5% BUSCO genes (Table 1). When compared to a k-mer database of the Hi-C reads, the pseudo-haplotypes range from 80.3% (pseudo-haplotype two from *L. fluviatilis*) to 89.7% (pseudo-haplotype one from *L. planeri*). The combined k-mer completeness was 92.2% for *L. fluviatilis* and 96.5% for *L. planeri* (Table 1). This completeness is visually represented in copy-number spectrum plots (Supplementary Figures 4-7). Overall, the consensus quality value (QV) of the different assemblies is high, from 28.9 (*L. planeri*, pseudo-haplotype two, compared to Hi-C k-mer database) to 56.0 (*L. fluviatilis*, pseudo-haplotype two, compared to the HiFi k-mer database). The QV is usually significantly higher when compared to the database of k-mers from the HiFi reads.

The Hi-C contact maps for the assemblies are shown in Supplementary Figure 8, and show a clear separation of the different chromosomes. GC-coverage plots for the assemblies are found in Supplementary Figure 9, showing similar coverage in the chromosomes with some spread in GC content.

For *L, fluviatilis, Flagger* identified 79.33% of pseudo-haplotype one as haploid, 20.35% as duplicated, 0.00% as error regions, and 0.03% as collapsed. The respective percentages for pseudo-haplotype two are 82.20% haploid, 17.54% duplicated, 0.0% error, and 0.03% collapsed (Table 1). For *L. planeri, Flagger* identified 75.27% of pseudo-haplotype one as haploid, 21.22% as duplicated, 3.20% as error regions, and 0.03% as collapsed. The respective percentages for pseudo-haplotype two are 75.57% haploid, 18.69% duplicated, 5.5% error, and 0.02% collapsed (Table 1).

We also aligned the pseudo-haplotypes of *L. fluviatilis* and *L. planeri* to each other and to another *L. fluviatilis* individual from the United Kingdom (kcLamFluv1; GCA_964198585.1) (Table 2 and Supplementary Table 2). The same settings did not give any results when used with *P. marinus*, it was likely too divergent from the *Lampetra* species.

**Table 2:** Different metrics based on alignment of pseudo-haplotype one of *L. fluviatilis* and *L. planeri* to each other and to an *L. fluviatilis* individual from the United Kingdom. See Supplementary Table 2 for metrics including pseudo-haplotype two.

| Aligned bases (percentage of genome assembly) |  |  |  |
| --- | --- | --- | --- |
|  | kcLamFluv2.2.h1 | kcLamPlan1.2.h1 | kcLamFluv1.1 |
| kcLamFluv2.2.h1 |  | 949,977,127<br>(90.5964%) | 927,780,578<br>(88.9979%) |
| kcLamPlan1.2.h1 | 958,093,979<br>(89.2691%) |  | 923,055,679<br>(88.5446%) |
| kcLamFluv1.1 | 952,584,705<br>(88.7558%) | 940,046,391<br>(89.6493%) |  |
| Insertions (sum in bp) |  |  |  |
|  | kcLamFluv2.2.h1 | kcLamPlan1.2.h1 | kcLamFluv1.1 |
| kcLamFluv2.2.h1 |  | 71,559 (179,071,556) | 68,502 (176,805,416) |
| kcLamPlan1.2.h1 | 74,939 (208,858,838) |  | 68,711 (179,916,524) |
| kcLamFluv1.1 | 77,251 (216,215,970) | 74,769 (190,651,971) |  |
| SNPs |  |  |  |
|  | kcLamFluv2.2.h1 | kcLamPlan1.2.h1 | kcLamFluv1.1 |
| kcLamFluv2.2.h1 |  | 4,547,241 | 4,604,025 |
| kcLamPlan1.2.h1 | 4,547,241 |  | 4,539,518 |
| kcLamFluv1.1 | 4,604,025 | 4,539,518 |  |
| Indels |  |  |  |
|  | kcLamFluv2.2.h1 | kcLamPlan1.2.h1 | kcLamFluv1.1 |
| kcLamFluv2.2.h1 |  | 5,879,800 | 5,949,776 |
| kcLamPlan1.2.h1 | 5,879,800 |  | 5,885,164 |
| kcLamFluv1.1 | 5,949,776 | 5,885,164 |  |

We also ran *OrthoFinder* on all the predicted proteins of the different assemblies and used *ASTRAL-Pro3* to generate quartet scores based on the gene trees from *OrthoFinder* (Figure 2). 40.1% of the gene trees placed *L. planeri* from Sweden (kcLamPlan1.2.h1) as a sister clade to *L. fluviatilis* from UK (kcLamFluv1.1) and *L. fluviatilis* from Norway (kcLamFluv2.2.h1) as a sister clade to *P. marinus*. 35.2% of the gene trees supported the *P. marinus* as sister clade to *L. fluviatilis* from UK (kcLamFluv1.1) and *L. fluviatilis* from Norway (kcLamFluv2.2.h1) as a sister clade to *L. planeri* from Sweden (kcLamPlan1.2.h1). Finally, 24.7% of the gene trees supported the last possible tree topology: *L. planeri* from Sweden (kcLamPlan1.2.h1) as a sister clade to *P. marinus* and *L. fluviatilis* from Norway (kcLamFluv2.2.h1) as a sister clade to *L. fluviatilis* from UK (kcLamFluv1.1).

**Figure 2:**
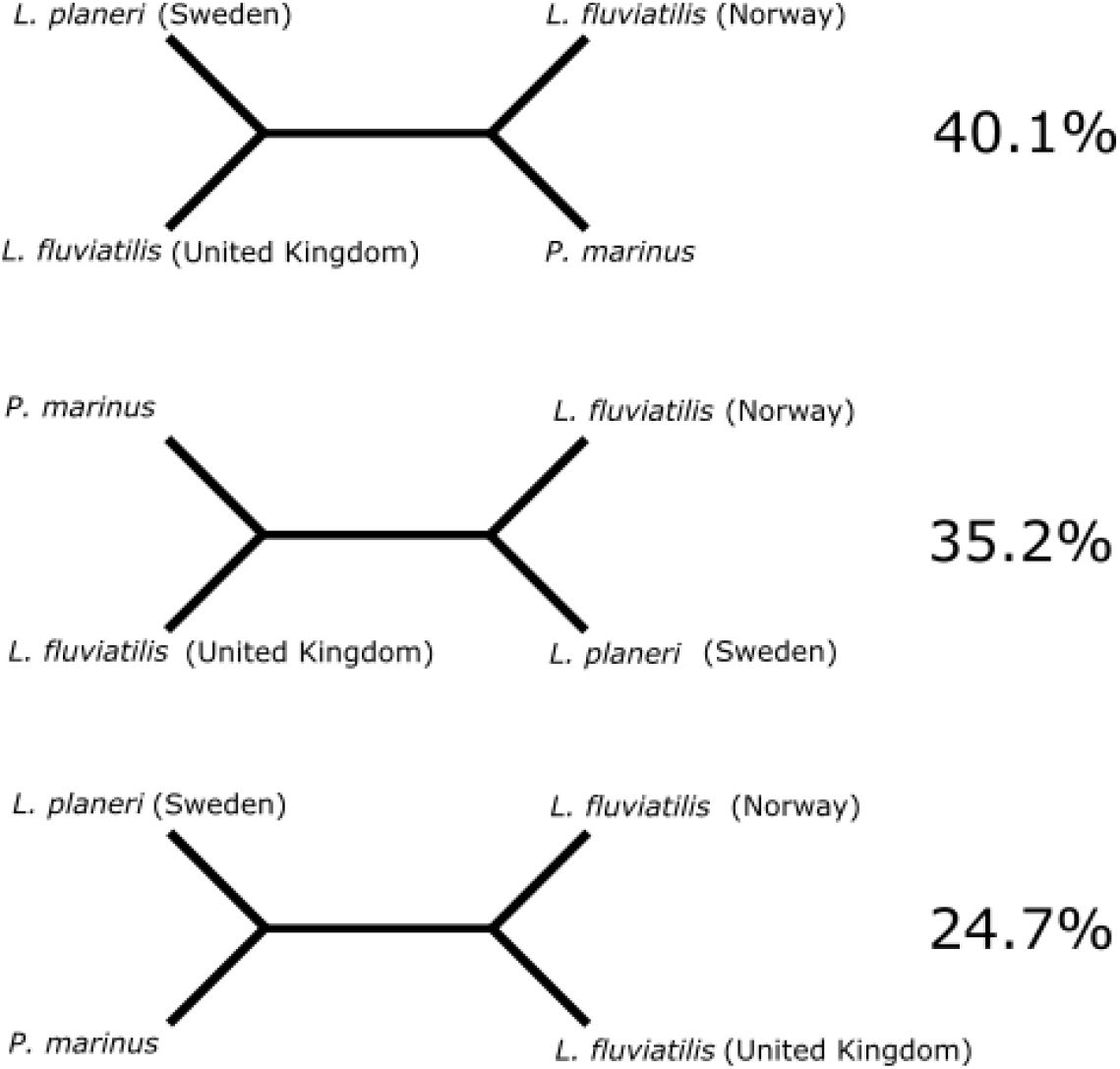
Different tree topologies and their support. The most complete pseudo-haplotype of *L. planeri* (hap1; called *L. planeri* (Sweden) in the figure) and of *L. fluviatilis* (hap1; called *L. fluviatilis* (Norway) in the figure) were used and compared with *L. fluviatilis* (UK) (kcLamFluv1; GCA_964198585.1) and *P. marinus* (kPetMar1; GCA_010993605.1). *ASTRAL-Pro3* was used to infer the species tree based on all gene trees from *OrthoFinder* and, in addition, to calculate the different quartet scores.

Gene order comparisons between the three different *Lampetra* individuals revealed conserved synteny among the genomes, with few chromosomal rearrangements (Figure 3). An increased number of reorganizations were observed when compared with the more distantly related *P. marinus* (Figure 3 and Supplementary Figure 10). In particular, chromosome 1 among the *Lampetra* individuals seems to be homologous across their length, while when compared to *P. marinus*, they are homologous to *P. marinus* chromosome 2 and chromosome 26. The same pattern can be observed with chromosome 2 among the *Lampetra* individuals, which were found to be homologous to chromosome 4 and chromosome 27 in comparison to *P. marinus* (Supplementary Figure 10). Moreover, chromosome 1 in the *Lampetra* individuals displays substantial connections between chromosome 1 as well as chromosome 10 when compared to other *Lampetra* individuals, indicating that these share shorter syntenic blocks along their length (Figure 3 and Supplementary Figure 10). This is also the case with chromosome 2 (homologous to chromosome 1 in *Lampetra*) and chromosome 10 in *P. marinus*.

**Figure 3:**
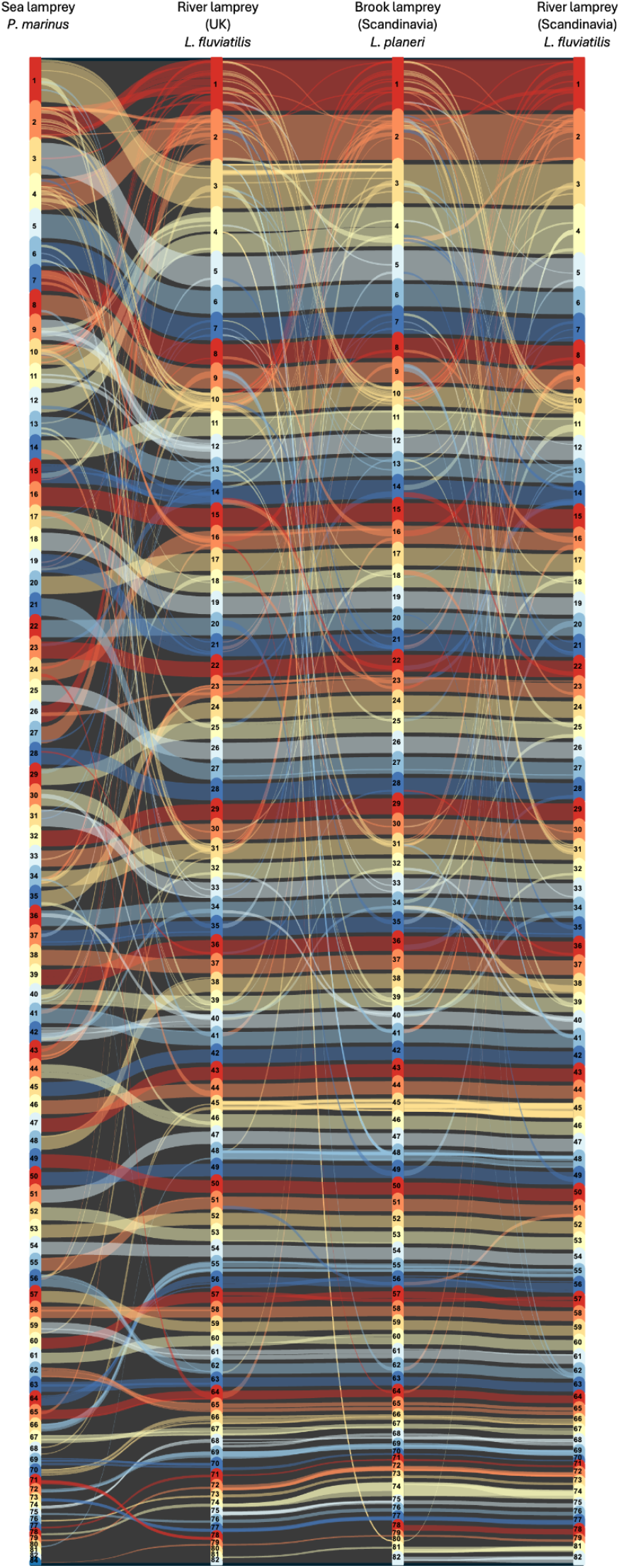
Chromosomal synteny between sea lamprey (*P. marinus*), river lamprey (UK; *L. fluviatilis*), brook lamprey (Scandinavia; *L. planeri*) and river lamprey (Scandinavia; *L. fluviatilis*). Chromosomal synteny of the most complete pseudo-haplotype of kcLamPlan1 and kcLamFluv2 (hap1 in both cases), kcLamFluv1 and kPetMar. Plots generated by *MCScanX* and *SynVisio* include chromosomes 1-82 for *L. fluviatilis* individuals and *L. planeri*, and 1-84 for *P. marinus*. Syntenic blocks are visualized as connected ribbons between individuals.

## Discussion

Here, we have sequenced, assembled, and annotated chromosome-level genomes from *L. planeri* and *L. fluviatilis*, resulting in two pseudo-haplotype separated assemblies. The reasons these assemblies differ in length could be due to heterogametic sex chromosomes/size differences in sex loci or some hitherto unknown chromosome diminishing (Marlétaz et al., 2024) affecting only one of the pseudo-haplotypes. It may also be due to unknown technical issues - more investigations are needed to resolve this. The pseudo-haplotype assemblies have comparable N50 statistics for both contigs (2.7-3.0 Mb here vs. 1.3 Mb for kcLamFluv1 and 2.5 Mb for kPetMar1). The scaffolds also had comparable N50 values (all around 13 Mb) as the previously released lamprey genome assemblies (kcLamFluv1 and kPetMar1) (Table 1 and Supplementary Figure 2). With regards to BUSCO scores, these are also comparable with 91.4% complete *BUSCO* genes in hap1 for *L. fluvialitis* (83.9% in hap2), 89.8% complete in hap1 for *L. planeri* (83.5% in hap2) and 91.8% in kcLamFluv1 and 92.5% in kPetMar1 (Table 1 and Supplementary Figure 2).

*Flagger* indicates that around 20% of the assemblies are duplicated. The *BUSCO* results do not support this (around 2-3% duplicated genes), however, we used the Metazoa marker gene set, with only 954 genes which could be too few to discover a putative duplication (Table 1). *GenomeScope* also only estimates 720-740 Mb genome sizes (about 20% less than the final assemblies) (Supplementary Figures 1 and 2). The common ancestor of lampreys and hagfish likely went through a triplication event of its genome (Yu et al., 2024), and this is likely reflected in the *Flagger* statistics and *GenomeScope* output as well as in the synteny plots. Interestingly, chromosomes 2 and 10 in *P. marinus* (1 and 10 in *Lampetra*) contain the two (of six in total) Hox clusters which do not have a clear ortholog relationship to the Hox clusters found in jawed vertebrates (Marlétaz et al., 2024). Our synteny analysis shows that there is collinearity between these chromosomes also in the *Lampetra* individuals, which shows that the pattern extends to multiple lamprey species (Figure 3).

If *L. fluviatilis* and *L. planeri* were two clearly differentiated species, we would expect more differences between them than between the two *L. fluviatilis* specimens. Based on the alignments between the different *Lampetra* individuals (Table 2), there is no clear separation between the two species. Rather, there are more differences between the two *L. fluviatilis* individuals with regards to indels and SNPs, than between either of the *L. fluviatilis* individuals and *L. planeri*. In contrast, there is no clear structure from insertions, depending on which assembly is query and target. Further, the largest fraction of the gene trees (40.1%) support *L. fluviatilis* (UK) and *L. planeri* as phylogenetic sister species, while only 24.7% support the two *L. fluviatilis* as sister species (Figure 2).

With regards to synteny there are only minor differences between the three *Lampetra* individuals - representing two *L. fluvatilis* from Norway and UK, respectively and an *L. planeri* from Sweden (fairly close to Norway: see details in Methods). With regards to chromosomal architecture, the results show that the genomes display conserved synteny with a few large rearrangements (Figure 3). The rearrangements that have taken place, when comparing the *Lampetra* individuals to the *P. marinus*, particularly involve chromosomes 1 and 2 (in *Lampetra*), which could be the result of lineage-specific fusions or fissions. Most notably, the differences in chromosomal architecture between *L. fluvatilis* and *L. planeri* are small compared to the geographical separation (Norway/Sweden and UK).

This study, based on two new high-quality reference genomes (*L. planeri* and *L. fluvatilis*) and a comparison with an *L. fluvatilis* reference genome from the UK may suggest that these represent a species complex with two ecotypes rather than two separate species. Thus, *L. planeri* and *L. fluvatilis* may represent two distinct possible life history trajectories of the same species. However, our study is only represented by 4 individuals (including the *P. marinus* outgroup individual). Even though the *L. fluviatilis* individual from Scandinavia robustly looks as different from *L. planeri* from Scandinavia as *L. fluviatilis* from the UK, the ultimate test for this conclusion would be to include whole genome sequenced individuals from multiple geographical locations across Europe - from the Mediterranean/South Atlantic oceans to the northern Atlantic. Ideally, such a study should also include spawning individuals to properly untangle the question of how the two putative ecotypes relate to each other.

## Acknowledgements

This project received data management and infrastructure support from ELIXIR Norway, supported by the Research Council of Norway grant 270068, the University of Bergen, the University of Oslo, the Arctic University of Norway in Tromsø, the Norwegian University of Science and Technology and the Norwegian University of Life Sciences: NMBU. The authors acknowledge support from the National Infrastructure for High Performance Computing and resources provided by Sigma2 as well as Data Storage in Norway (project NN8013K) for computational work. The Norwegian Sequencing Centre generated the sequencing data used in this project (http://sequencing.uio.no). The authors especially thank Sarah Fullmer for careful reading and feedback on the text.

## Data Availability

Data generated for this study are available under ENA BioProject PRJEB77187 and PRJEB77192 for EBP-Nor. Raw PacBio sequencing data for *L. fluviatilis* (ENA BioSample: SAMEA115797768) are deposited in ENA under ERX12712303, ERX12712308 and ERX12712309, while Illumina Hi-C sequencing data is deposited in ENA under ERX12712501. Pseudo-haplotype one can be found in ENA at PRJEB77117 while pseudo-haplotype two is PRJEB77186. Raw PacBio sequencing data for *L. planeri* (ENA BioSample: SAMEA115802553) are deposited in ENA under ERX12713780, ERX12713797 and ERX12713807, while Illumina Hi-C sequencing data is deposited in ENA under ERX12714064. Pseudo-haplotype one can be found in ENA at PRJEB77190 while pseudo-haplotype two is PRJEB77191.

The genome annotations are available at https://zenodo.org/records/14288109 (DOI:10.5281/zenodo.11159637).

## Authors’ contributions’

**Ole K. Tørresen:** Writing - original draft, Formal analysis, Visualization, Writing - review and editing. **Benedicte Garmann-Aarhus:** Writing - original draft, Investigation, Formal analysis, Visualization. **Siv Nam Khang Hoff:** Writing - original draft, Formal analysis, Visualization. **Sissel Jentoft:** Writing - review and editing. **Mikael Svensson:** Resources. **Eivind Schartum:** Resources. **Ave Tooming-Klunderud:** Investigation. **Morten Skage:** Investigation. **Anders Krabberød:** Formal analysis. **Leif Asbjørn Vøllestad:** Project Writing - original draft, Writing - review and editing, administration. **Kjetill S. Jakobsen:** Project administration, Writing - original draft, Writing - review and editing, Funding acquisition.

