## Supplementary information for "Comparison of whole-genome assemblies of European river lamprey (*Lampetra fluviatilis*) and brook lamprey (*Lampetra planeri*)"

**Supplementary figures:**

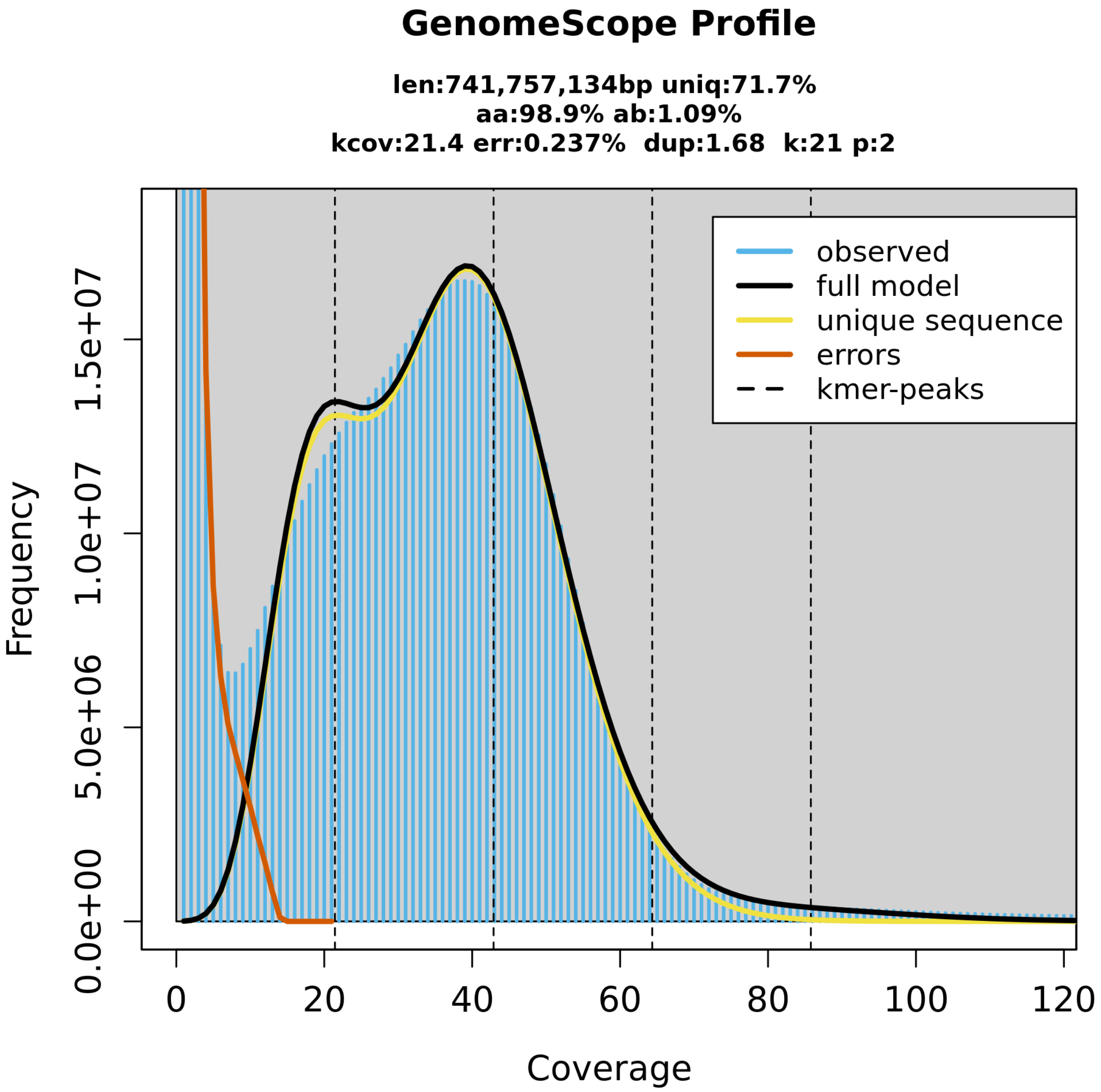


**Supplementary Figure 1: Genome profile of *L. fluviatilis*.**. This analysis estimates a 742 Mb genome, with 1.09 % heterozygosity and bimodal pattern characteristic of a diploid genome. While the peaks of the k-mer plot are not clearly visible, there is a left shoulder which corresponds to k-mers from heterozygous regions of the genome, while the right-hand peak is from homozygous regions.

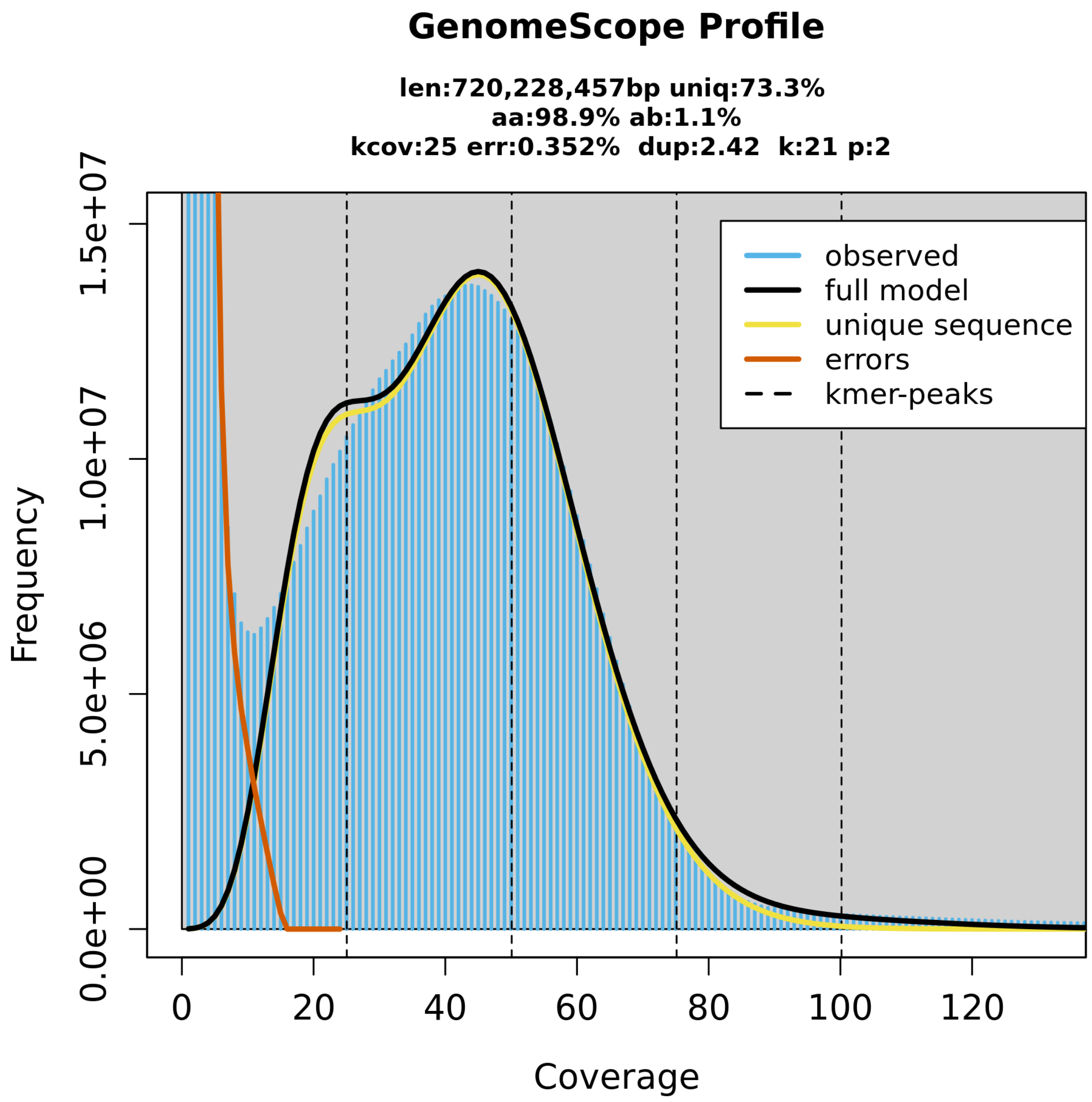


**Supplementary Figure 2: Genome profile of *L. planeri*.**. This analysis estimates a 720 Mb genome, with 1.1 % heterozygosity and bimodal pattern characteristic of a diploid genome. While the peaks of the k-mer plot are not clearly visible, there is a left shoulder which corresponds to k-mers from heterozygous regions of the genome, while the right-hand peak is from homozygous regions.

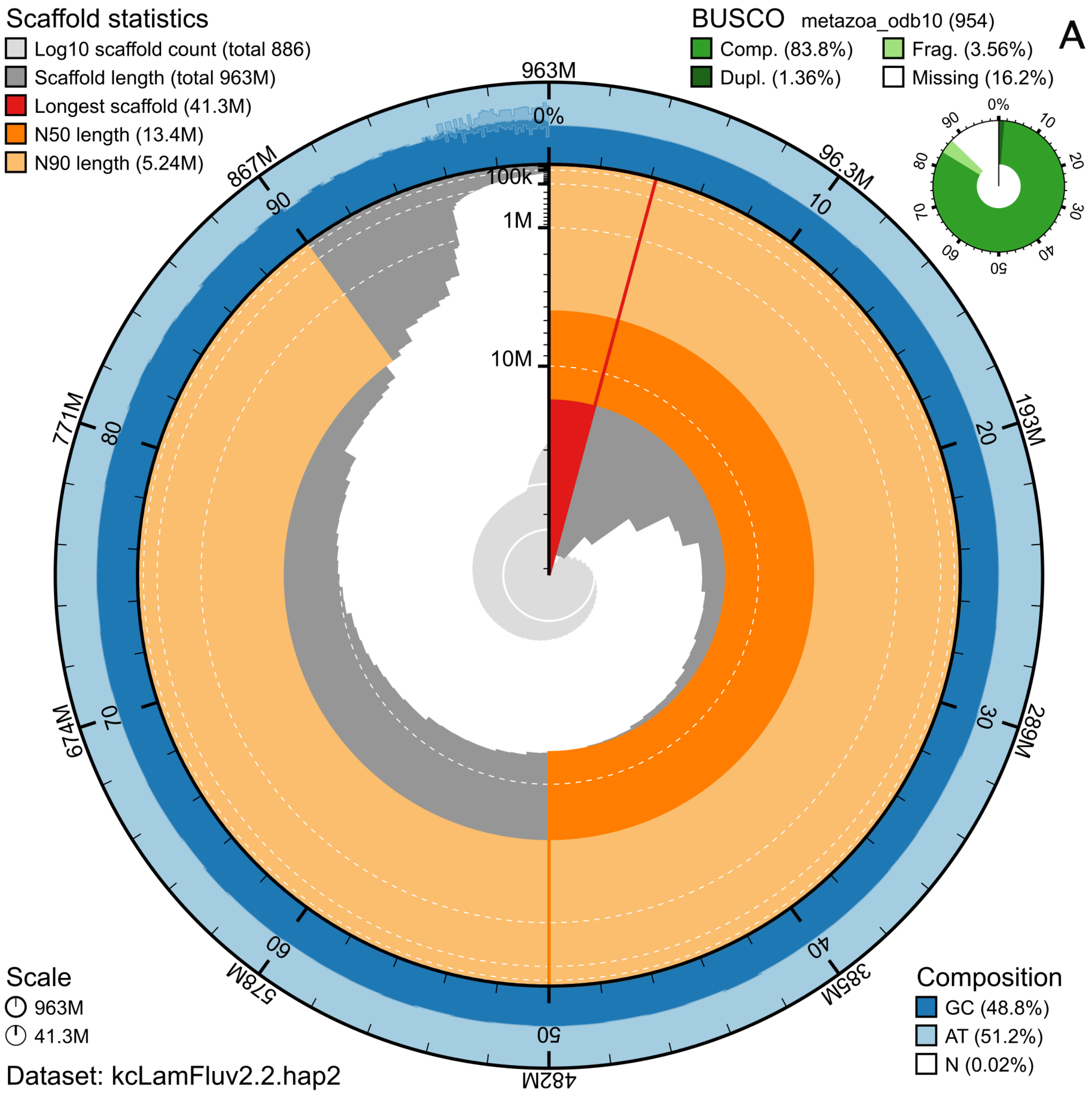

Scaffold statistics
Log10 scaffold count (total 886)
Scaffold length (total 963M)
Longest scaffold (41.3M)
N50 length (13.4M)
N90 length (5.24M)
BUSCO metazoa_odb10 (954)
Comp. (83.8%)
Dupl. (1.36%)
Frag. (3.56%)
Missing (16.2%)
A
963M
96.3M
193M
289M
385M
482M
578M
674M
771M
867M
0%
10
20
30
40
50
60
70
80
90
100k
1M
10M
Scale
963M
41.3M
Dataset: kcLamFluv2.2.hap2
Composition
GC (48.8%)
AT (51.2%)
N (0.02%)

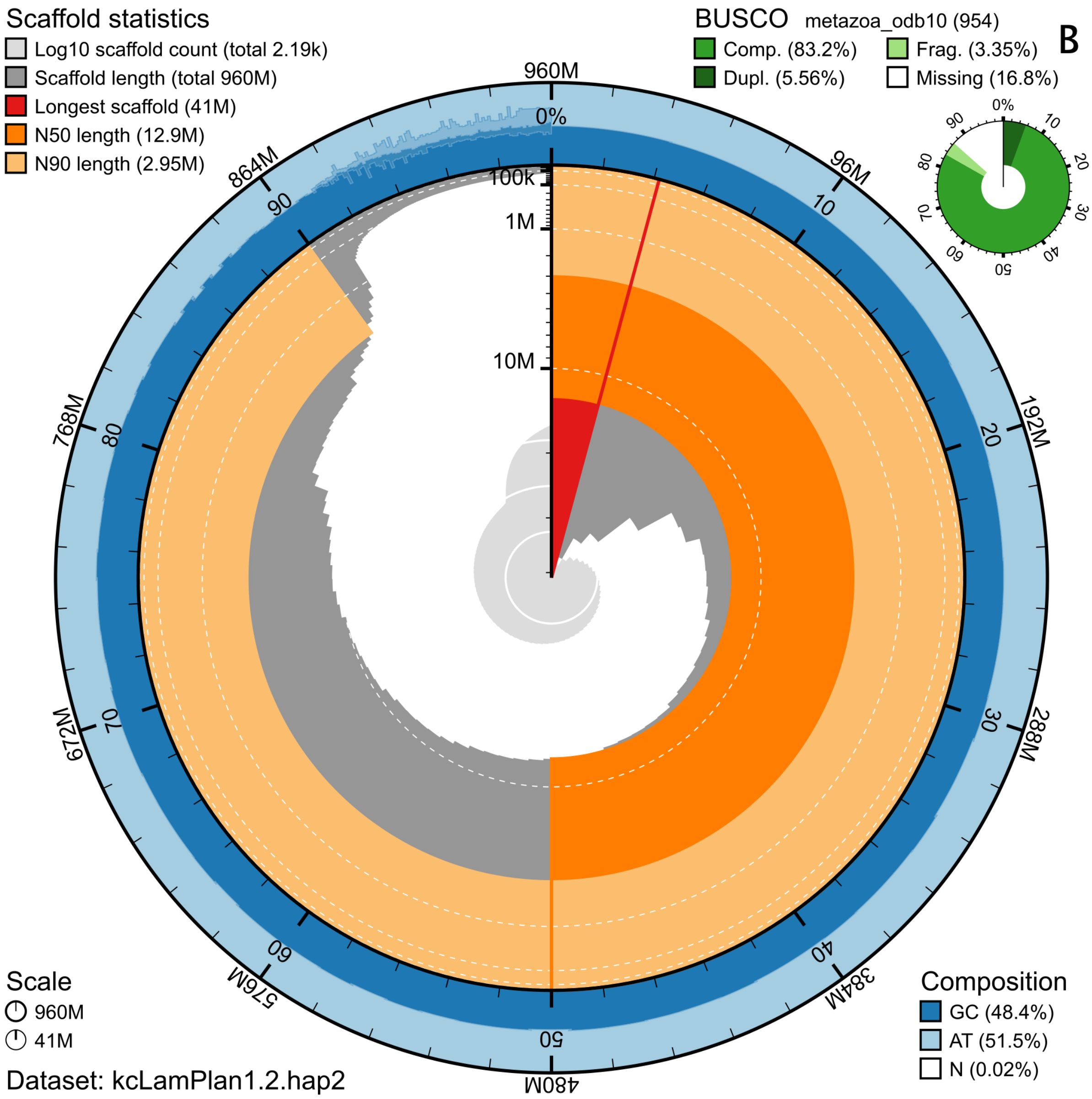


**Supplementary Figure 3: Metrics of the genome assemblies of *L. fluviatilis* (A) and *L. planeri* (B), pseudo-haplotype two for both species.** The BlobToolKit Snailplots show N50 metrics and BUSCO gene completeness. The two outermost bands of the circle signify GC versus AT composition at 0.1% intervals, with mean, maximum and minimum. Light orange shows the N90 scaffold length, while the deeper orange is N50 scaffold length. The red line shows the size of the largest scaffold. All the scaffolds are arranged in a clockwise manner from the largest to the smallest and are shown in darker gray with white lines at different orders of magnitude, while the light gray shows cumulative count of scaffolds.

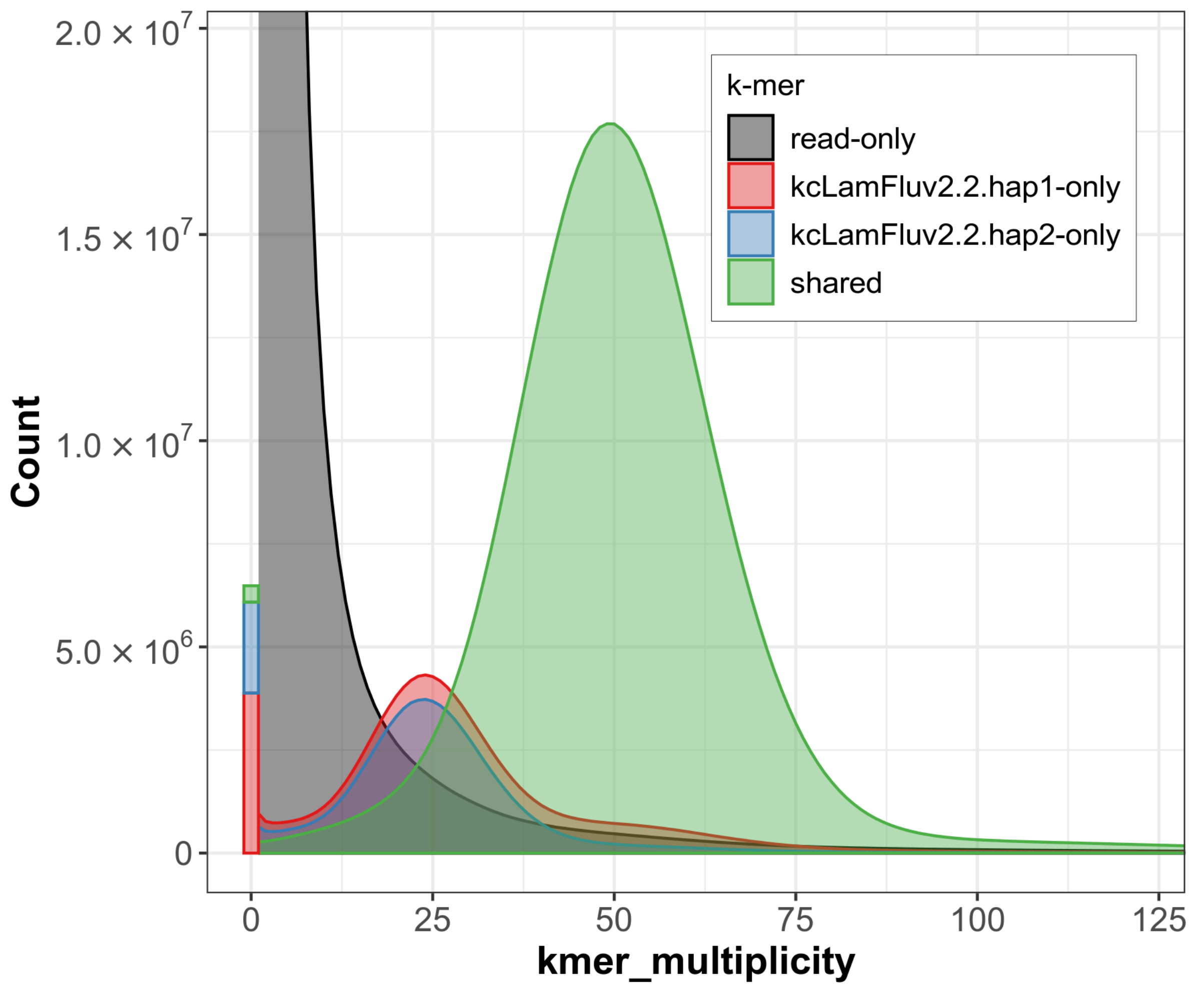


**Supplementary Figure 4: K-mer copy-number spectrum analysis of *L. fluviatilis* compared to k-mers from a database from the Hi-C reads.** Assembly-specific k-mers are shown in red and blue, while k-mers shared by both pseudo-haplotypes are in green. The stack above 0 on the x-axis shows k-mers found in the assemblies, but not in the reads. Figure is generated by Merqury.

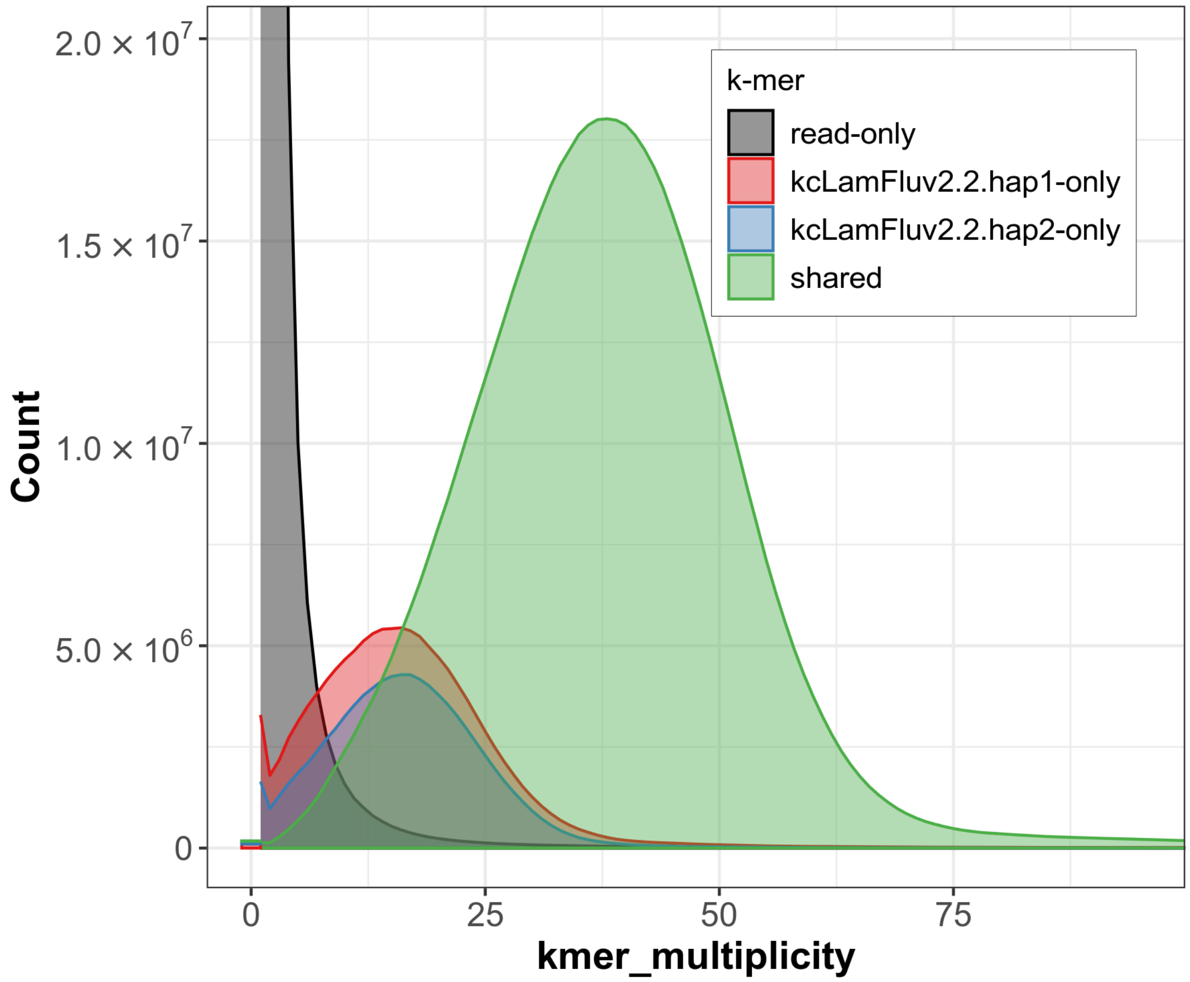


**Supplementary Figure 5: K-mer copy-number spectrum analysis of *L. fluviatilis* compared to k-mers from a database from the PacBio HiFi reads.** Assembly-specific k-mers are shown in red and blue, while k-mers shared by both pseudo-haplotypes are in green. The stack above 0 on the x-axis shows k-mers found in the assemblies, but not in the reads. Figure is generated by Merqury.

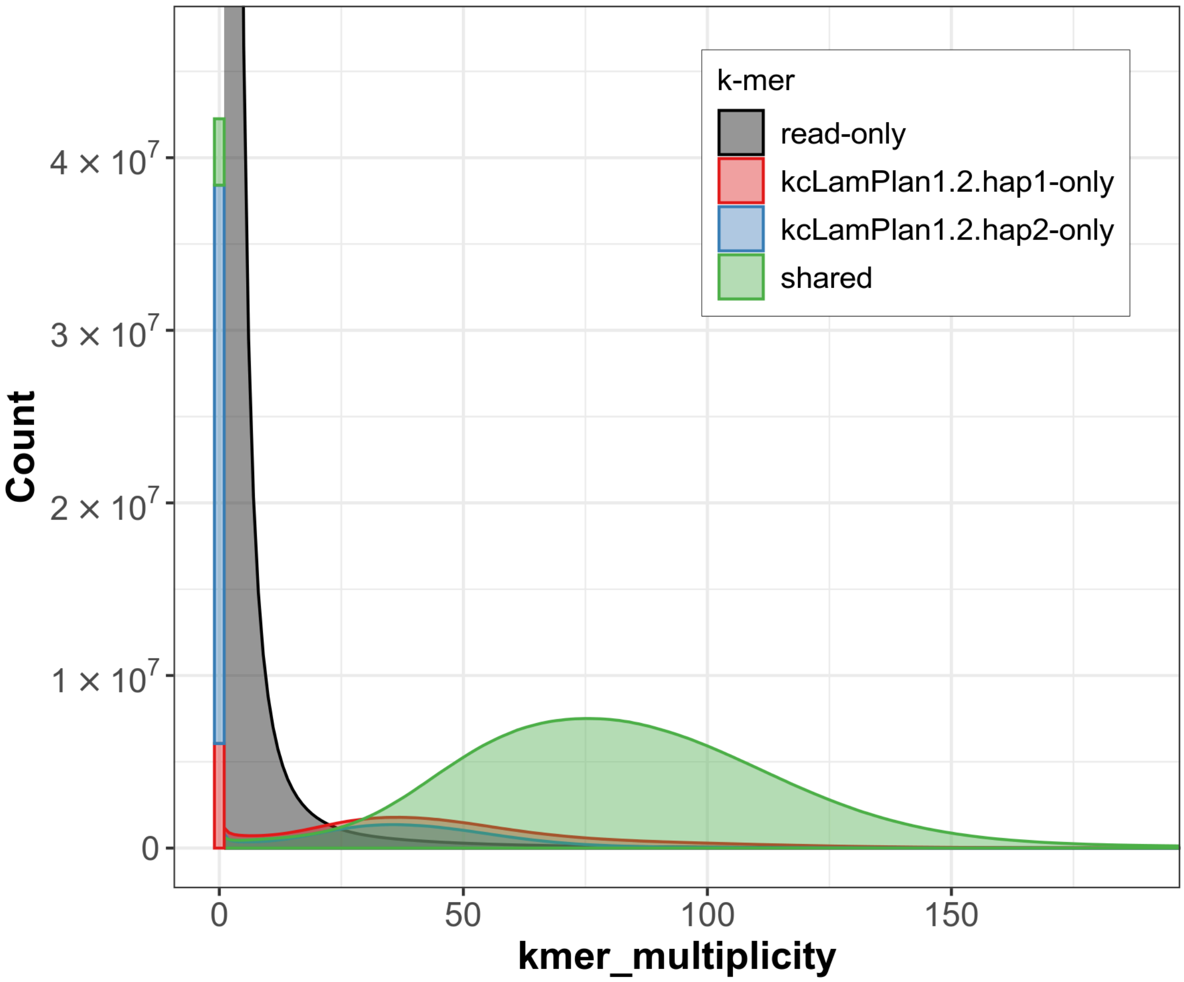


**Supplementary Figure 6: K-mer copy-number spectrum analysis of *L. planeri* compared to k-mers from a database from the Hi-C reads.** Assembly-specific k-mers are shown in red and blue, while k-mers shared by both pseudo-haplotypes are in green. The stack above 0 on the x-axis shows k-mers found in the assemblies, but not in the reads. Figure is generated by Merqury.

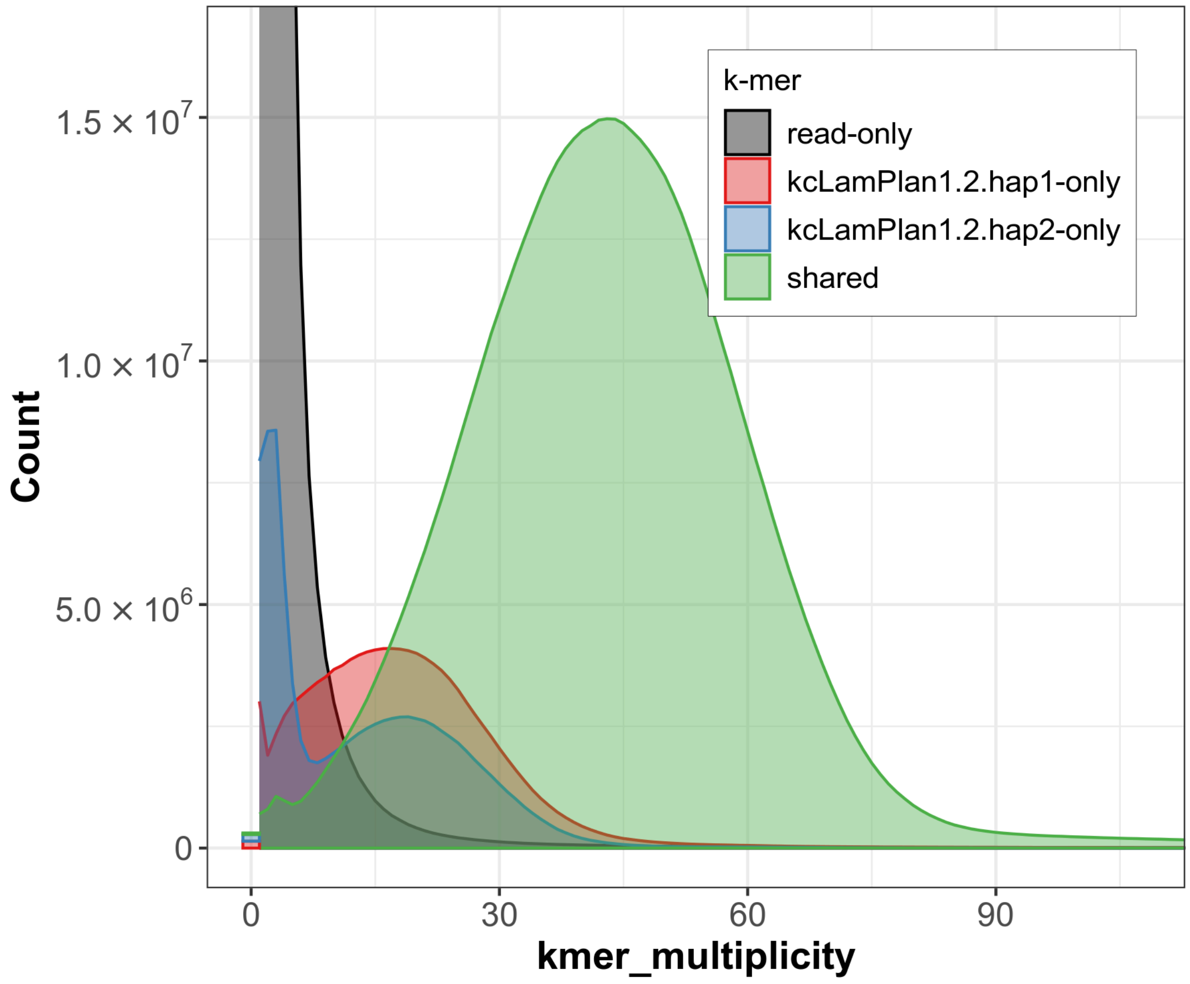


**Supplementary Figure 7: K-mer copy-number spectrum analysis of *L. planeri* compared to k-mers from a database from the PacBio HiFi reads.** Assembly-specific k-mers are shown in red and blue, while k-mers shared by both pseudo-haplotypes are in green. The stack above 0 on the x-axis shows k-mers found in the assemblies, but not in the reads. Figure is generated by Merqury.

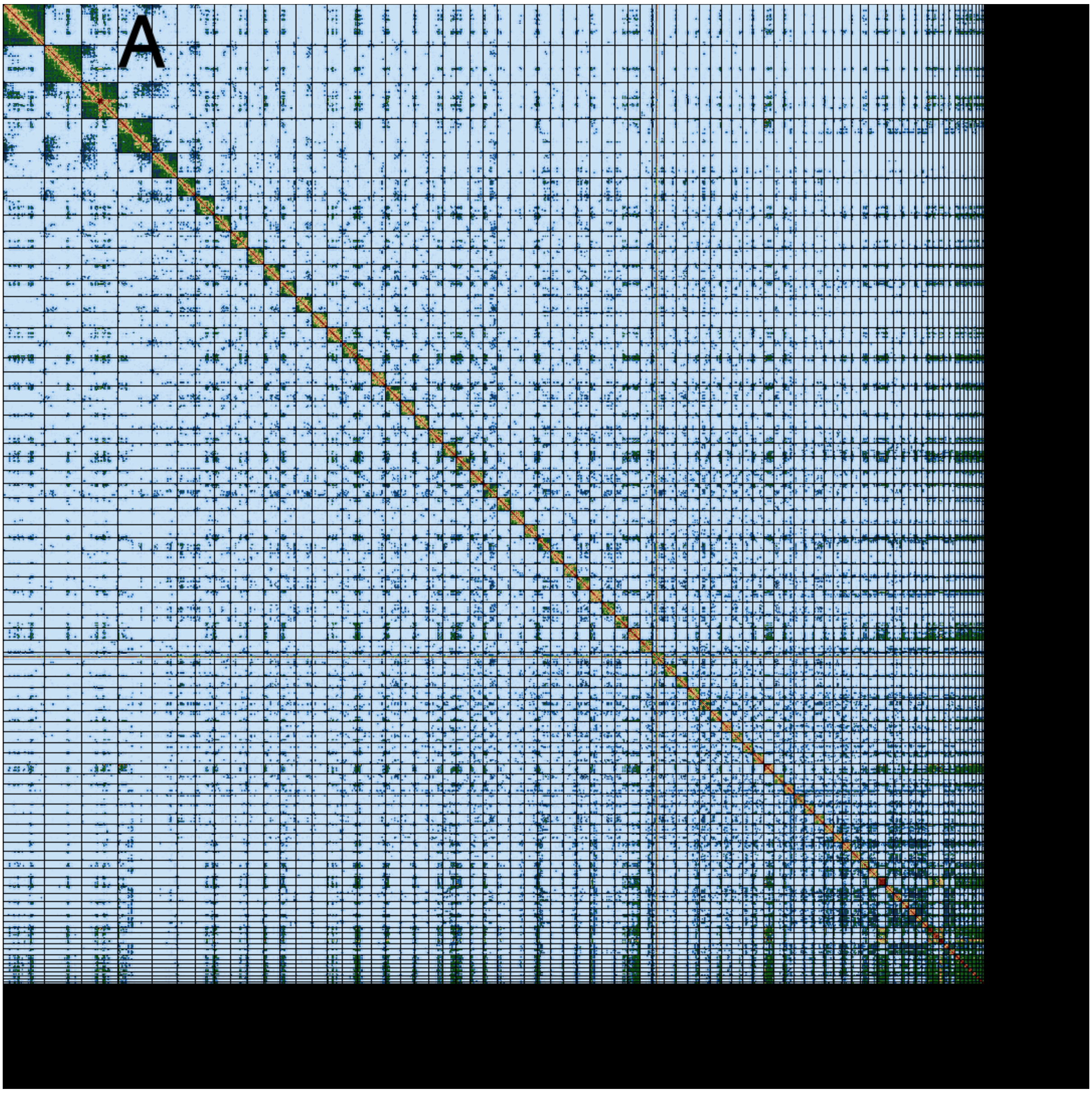
A

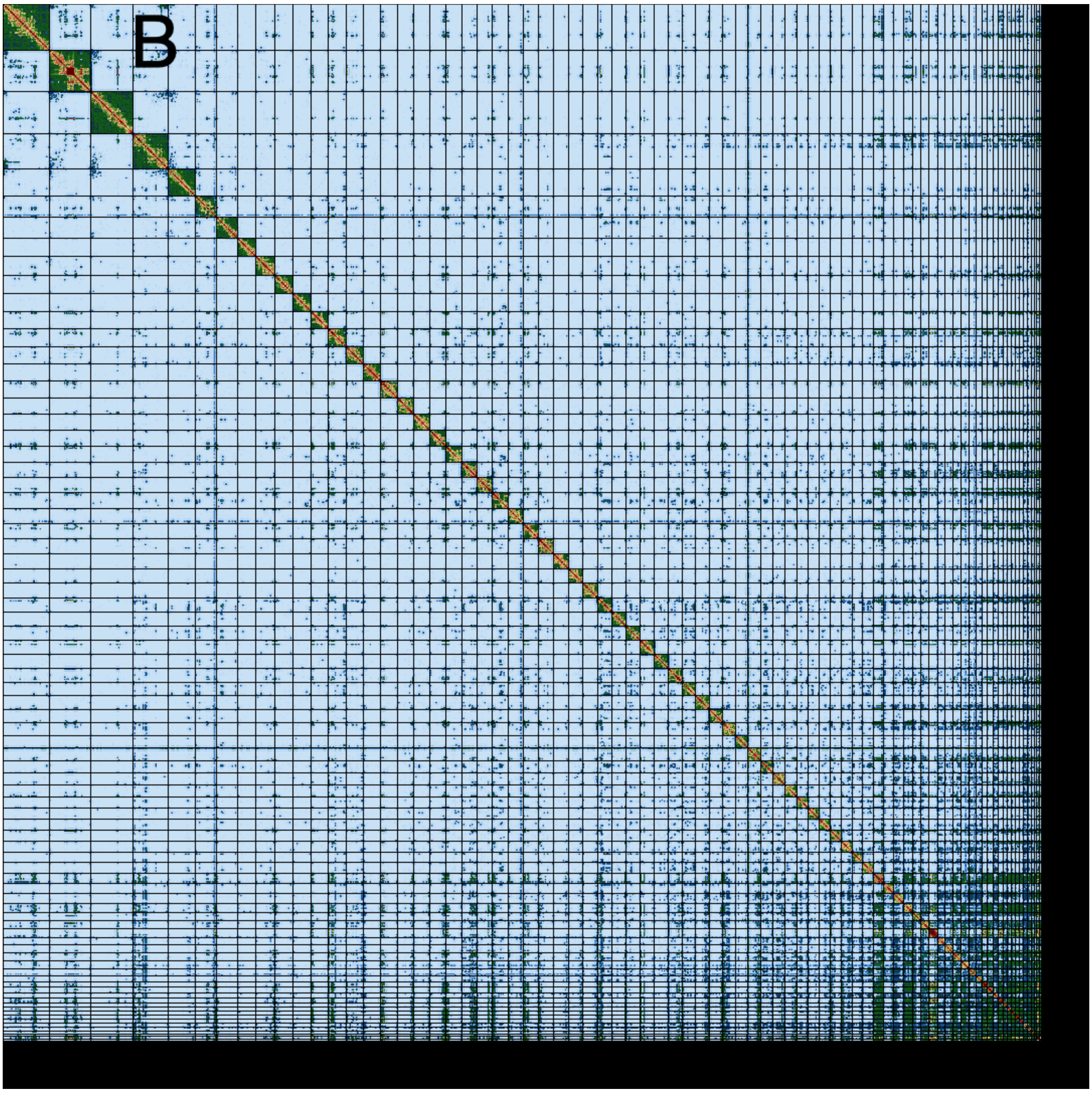
B

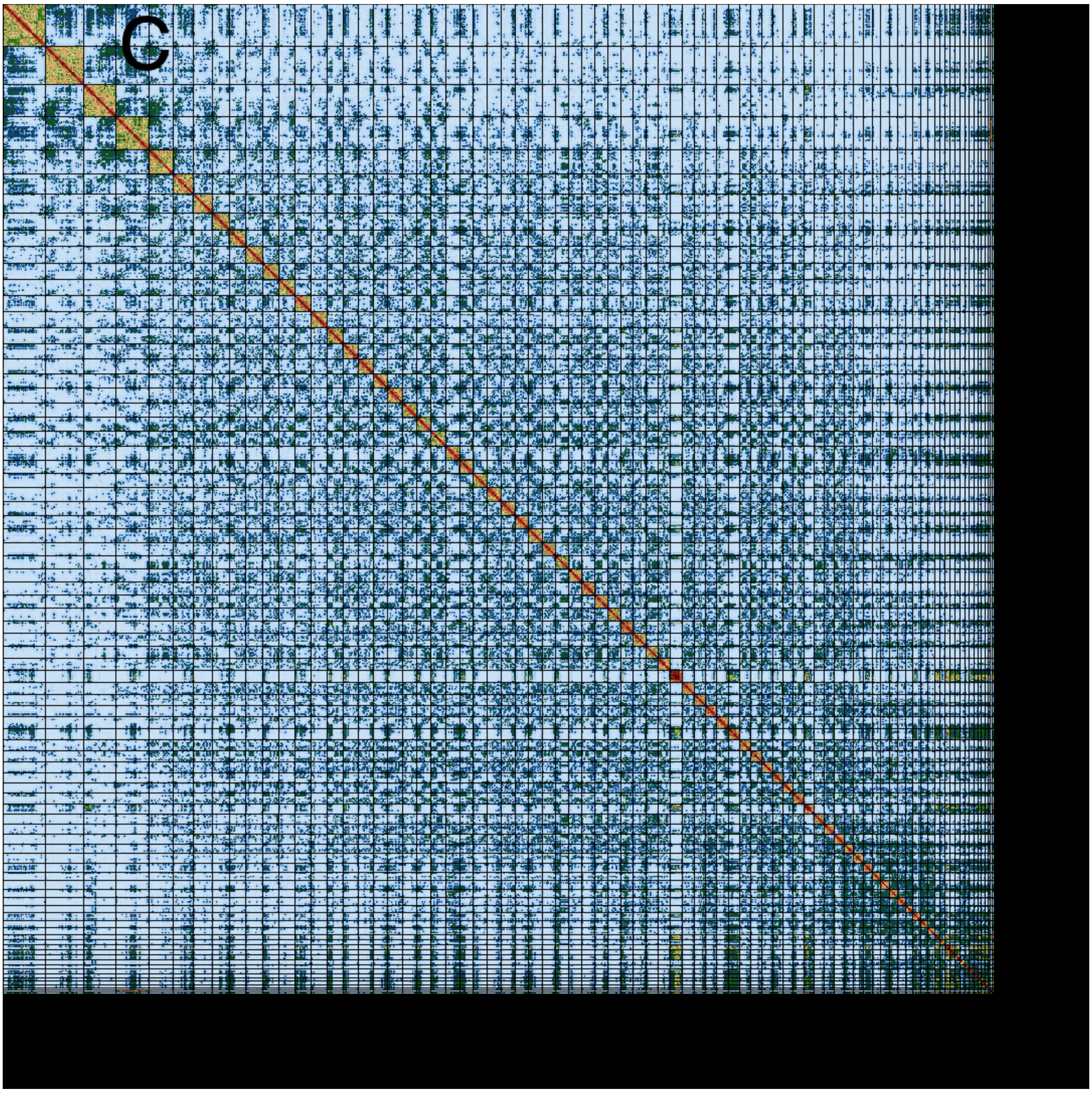
C

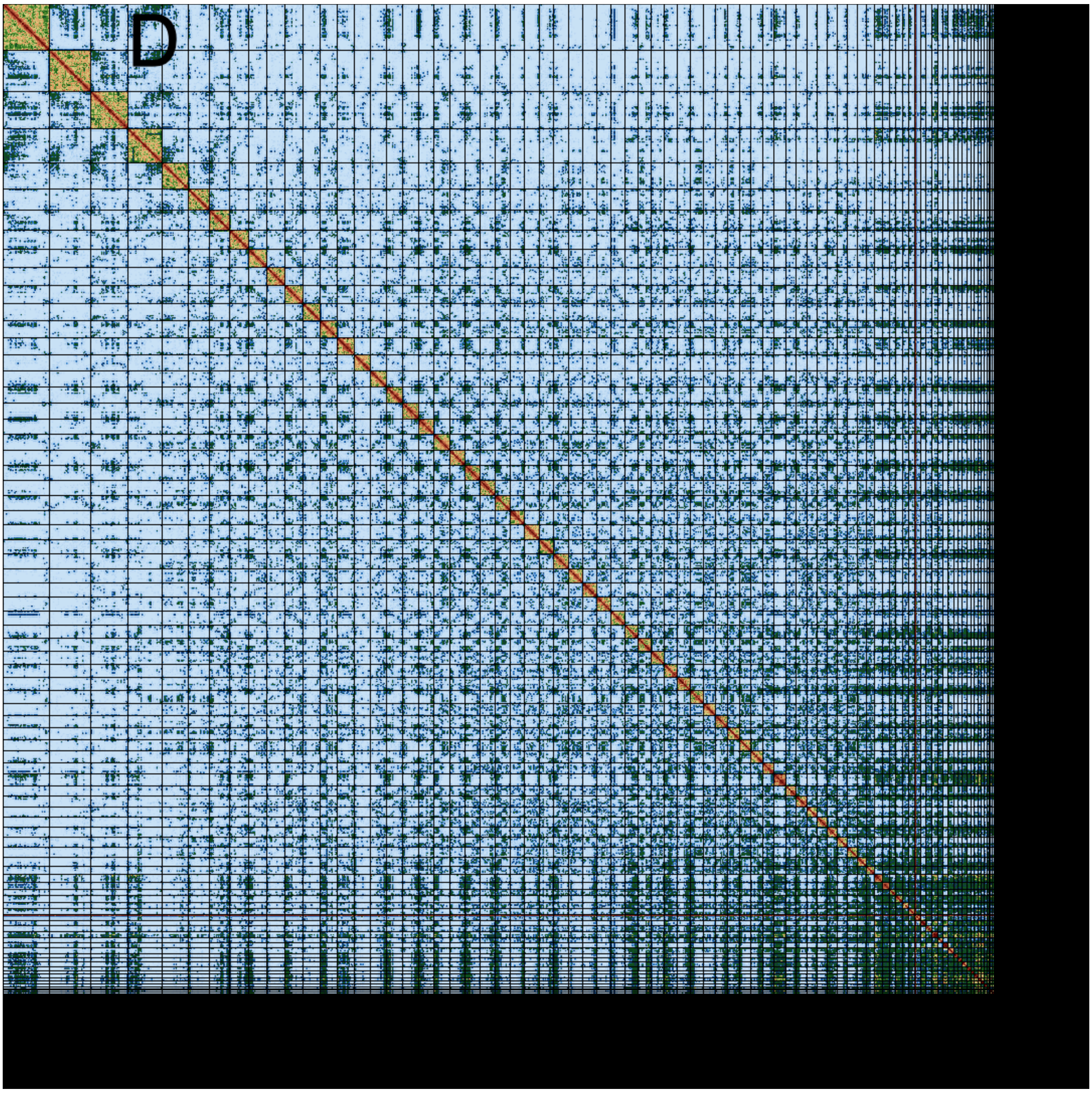


**Supplementary Figure 8: Hi-C contact map of genome assemblies of *L. fluviatilis* and *L.planeri* for hap1 and hap2 in both species.** All assemblies are visualized using PreTextSnapshot. Chromosomes are shown in order of size from left to right and top to bottom. A: kcLamFluv2.2.hap1, B: kcLamFluv2.2.hap2, C:kcLamPlan1.2.hap1 and D:kcLamPlan1.2.hap2. The black bands to the right and bottom consist of small contigs which the resolution is too low to distinguish.

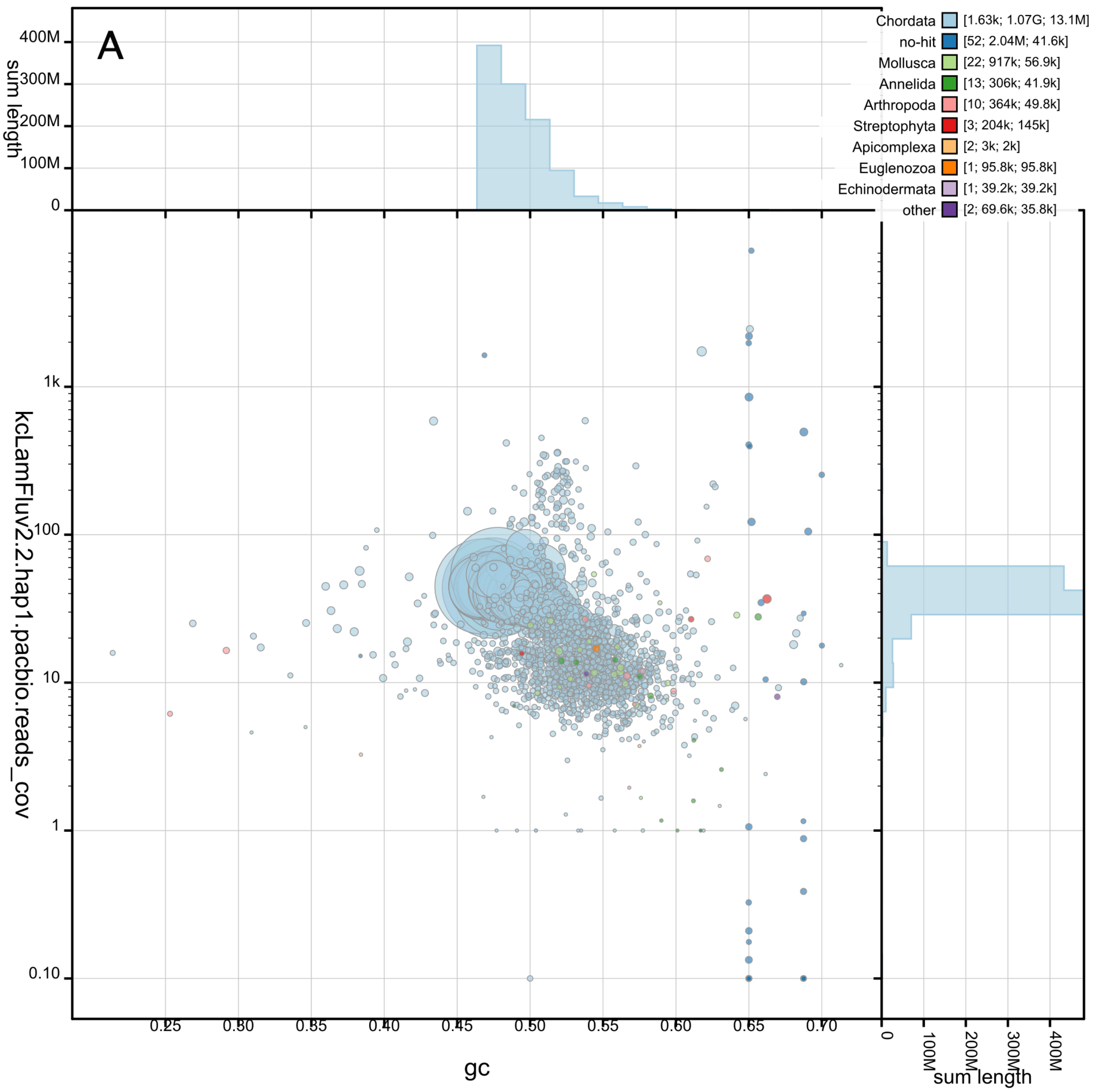

A
400M
300M
200M
100M
0
sum length
Chordata [1.63k; 1.07G; 13.1M]
no-hit [52; 2.04M; 41.6k]
Mollusca [22; 917k; 56.9k]
Annelida [13; 306k; 41.9k]
Arthropoda [10; 364k; 49.8k]
Streptophyta [3; 204k; 145k]
Apicomplexa [2; 3k; 2k]
Euglenozoa [1; 95.8k; 95.8k]
Echinodermata [1; 39.2k; 39.2k]
other [2; 69.6k; 35.8k]
kcLamFluv2.2.hap1.pacbio.reads_cov
1k
100
10
1
0.10
0.25
0.30
0.35
0.40
0.45
0.50
0.55
0.60
0.65
0.70
gc
0
100M
200M
300M
400M
sum length

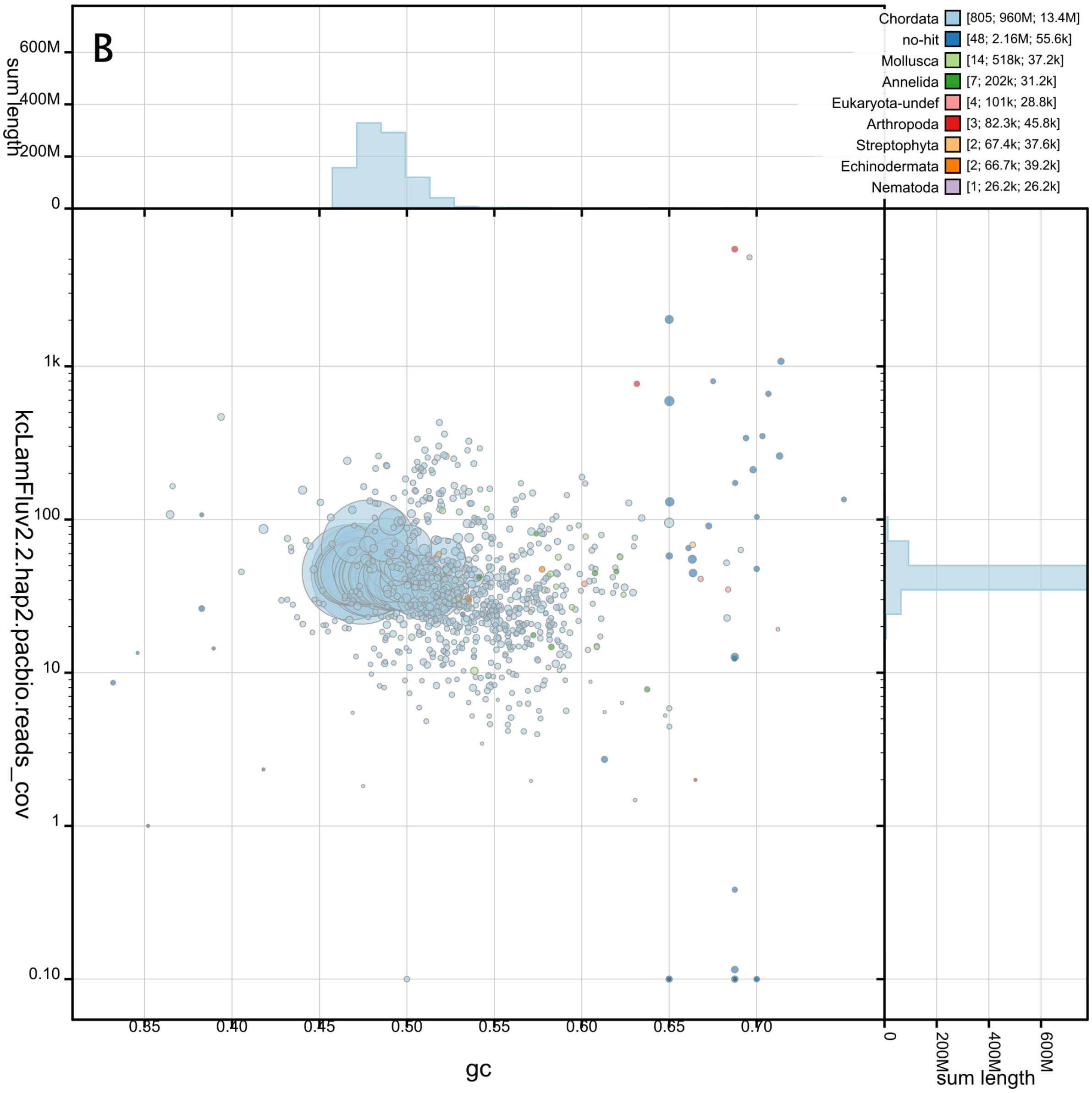
B
Chordata [805; 960M; 13.4M]
no-hit [48; 2.16M; 55.6k]
Mollusca [14; 518k; 37.2k]
Annelida [7; 202k; 31.2k]
Eukaryota-undef [4; 101k; 28.8k]
Arthropoda [3; 82.3k; 45.8k]
Streptophyta [2; 67.4k; 37.6k]
Echinodermata [2; 66.7k; 39.2k]
Nematoda [1; 26.2k; 26.2k]
sum length
600M
400M
200M
0
kcLamFluv2.2.hap2.pacbio.reads_cov
1k
100
10
1
0.10
0.35
0.40
0.45
0.50
0.55
0.60
0.65
0.70
gc

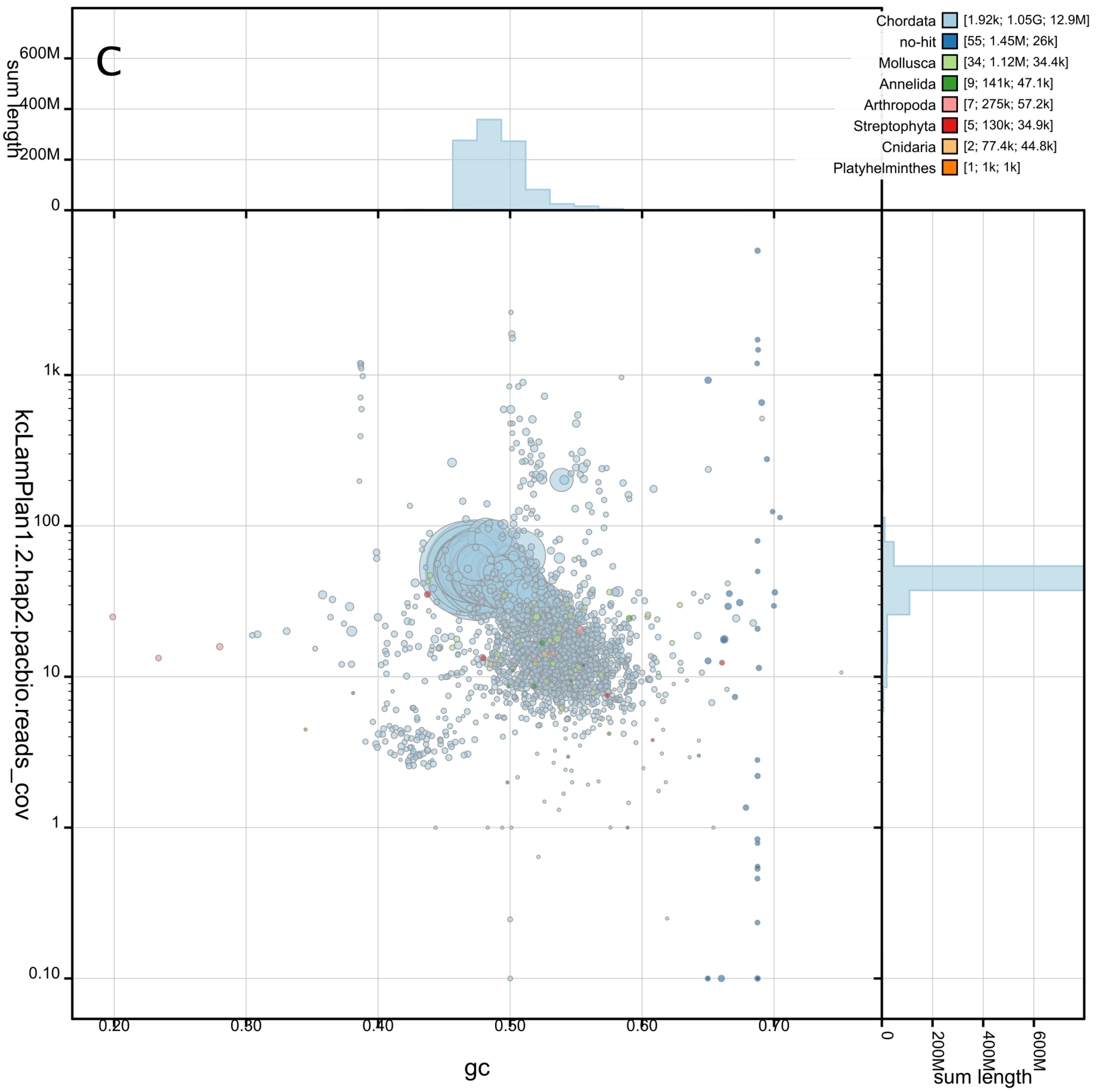

C
Chordata [1.92k; 1.05G; 12.9M]
no-hit [55; 1.45M; 26k]
Mollusca [34; 1.12M; 34.4k]
Annelida [9; 141k; 47.1k]
Arthropoda [7; 275k; 57.2k]
Streptophyta [5; 130k; 34.9k]
Cnidaria [2; 77.4k; 44.8k]
Platyhelminthes [1; 1k; 1k]
sum length
600M
400M
200M
0
kcLamPlan1.2.hap2.pacbio.reads_cov
1k
100
10
1
0.10
0.20
0.30
0.40
0.50
0.60
0.70
gc
0
200M
400M
600M
sum length

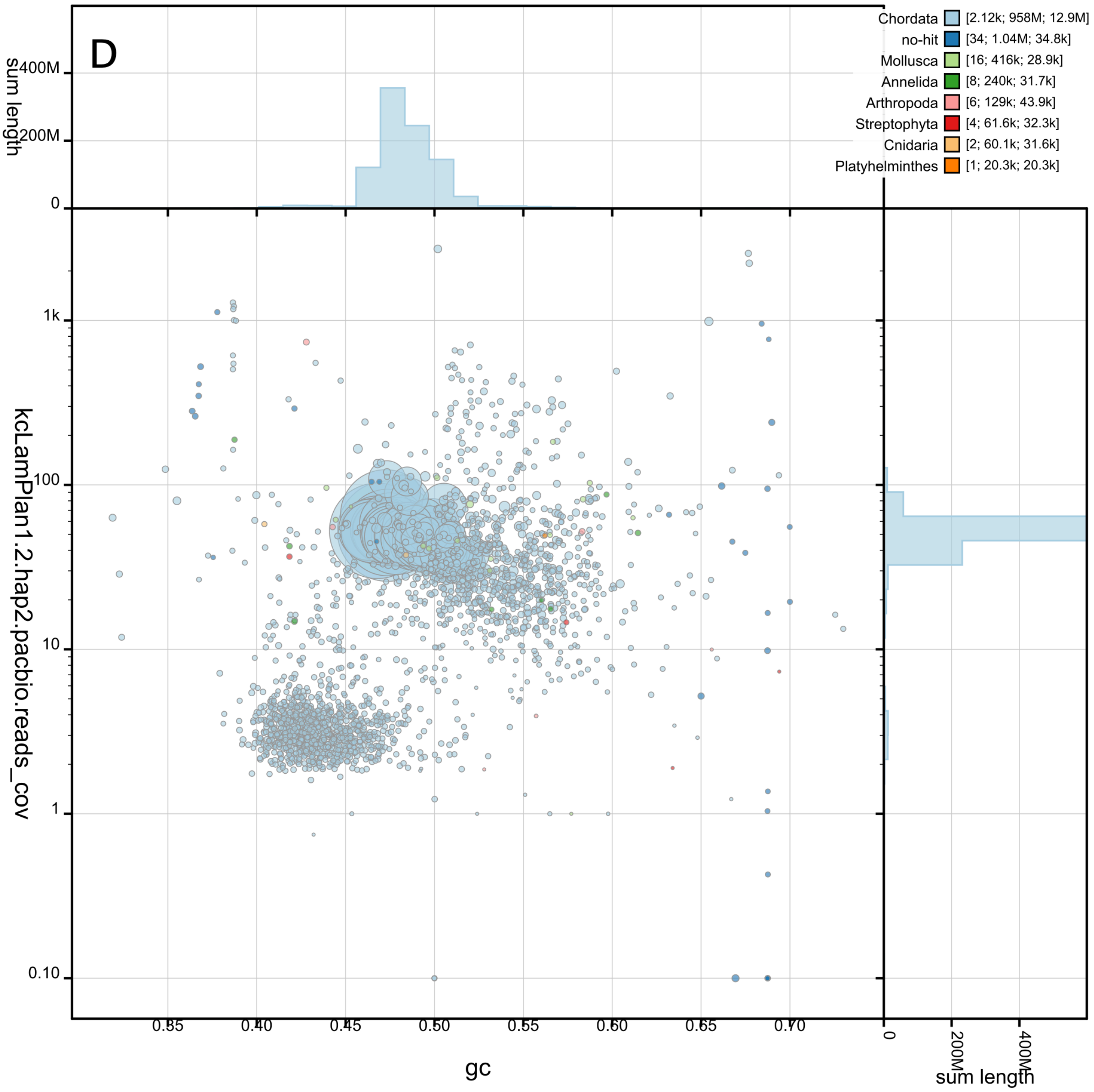


**Supplementary Figure 9: BlobToolKit GC-coverage plots of genome assemblies of *L. fluviatilis* hap1 (A) and hap2 (B), and *L. planeri* hap1 (C) and hap2 (D).** The scaffolds are coloured by phylum. The size of the circles are in proportion to the length of the scaffolds. Histograms show the distribution of scaffold length sum along each axis.

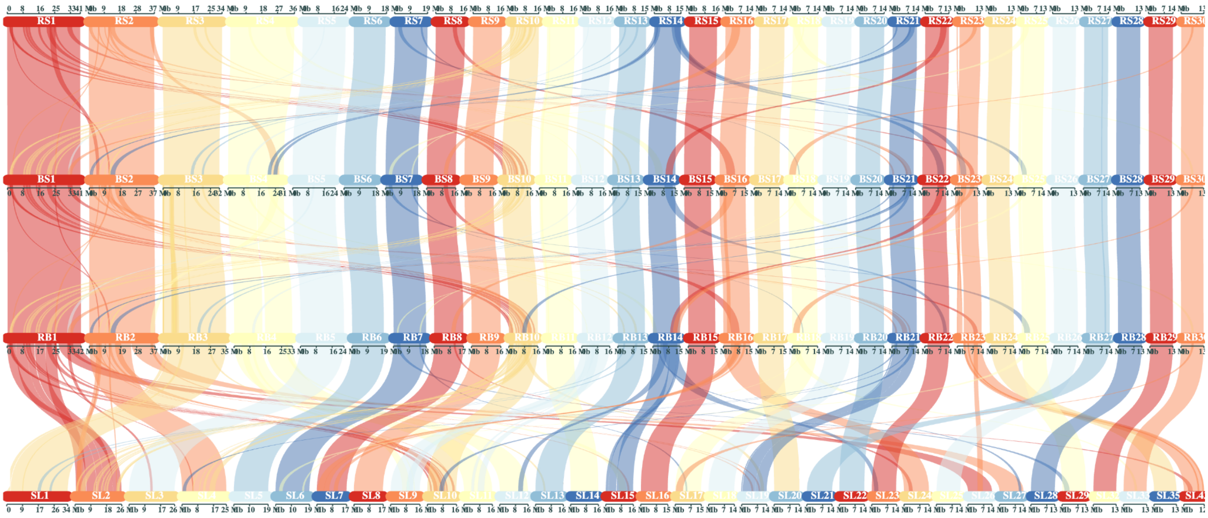

**Supplementary Figure 10: Synteny plot based on *MCScanX* and *Synvisio* for the 30 longest chromosomes of the most complete pseudo-haplotype of *L. planeri* (kcLamPlan1; BS [Brook Scandinavia]) and *L. fluviatilis* (kcLamFluv2; RS [River Scandinavia]) (hap1 in both cases), *L. fluviatilis* (kcLamFluv1; RB [River Britain]) as well as the homologous chromosomes in *P. marinus* (kPetMar; SL [Sea Lamprey]).** Syntenic blocks are visualized as connection ribbons between individuals. Chromosomal sizes are shown in a legend below each genome.

### Supplementary Tables:

**Supplementary Table 1. Software tools: versions and sources**

| Software tool | Version | Source |
|---|---|---|
| BlobToolKit | 4.1.7 | https://github.com/blobtoolkit/blobtoolkit |
| blobtk | 0.5.1 | https://github.com/blobtoolkit/blobtk |
| BUSCO | v5.4.7 | https://gitlab.com/ezlab/busco |
| hifiasm | 0.16.1-r375 | https://github.com/chhylp123/hifiasm |
| KMC | v3.1.2rc1 | https://github.com/refresh-bio/KMC |
| GenomeScope | v2.0 | https://github.com/tbenavi1/genomescope2.0 |
| Smudgeplot | 1.2.5 | https://github.com/KamilSJaron/smudgeplot |
| HiFiAdapterFilt | v2.0.0 | https://github.com/sheinasim/HiFiAdapterFilt |
| PretextView | 0.2.5 | https://github.com/wtsi-hpag/PretextView |
| PretextMap | 0.1.9 | https://github.com/wtsi-hpag/PretextMap |
| PretextSnapshot | commit 16b42f2 | https://github.com/wtsi-hpag/PretextSnapshot |
| bedtools | 2.30.0 | https://github.com/arq5x/bedtools2 |
| meryl | 1.3.0 | https://github.com/marbl/meryl |
| BWA-MEM | v0.7.17 | https://github.com/lh3/bwa |
| samtools | 1.17 | https://github.com/samtools/samtools |
| YaHS | yahs-1.1.91eebc2 | https://github.com/c-zhou/yahs |
| FCS-GX | 0.3.0 | https://github.com/ncbi/fcs |
| Merqury | v1.3 | https://github.com/marbl/merqury |
| AGAT | v1.0 | https://github.com/NBISweden/AGAT |
| MitoHiFi | v2.2 | https://github.com/marcelauliano/MitoHiFi |
| miniprot | 0.11-r234 | https://github.com/lh3/miniprot |
| GALBA | 1.0.6 | https://github.com/Gaius-Augustus/GALBA |
| RED | v2018.09.10 | http://toolsmith.ens.utulsa.edu/ |
| Funannotate | v1.8.13 | https://github.com/nextgenusfs/funannotate |
| EvidenceModeler | v1.1.1 | https://github.com/EVidenceModeler/EVidenceModeler |
| DIAMOND | v2.0.15<br>v2.1.6* | https://github.com/bbuchfink/diamond |

| | | |
|---|---|---|
| InterProScan | v5.47-82 | https://www.ebi.ac.uk/interpro/search/sequence/ |
| EMBLmyGFF3 | v2.2 | https://github.com/NBISweden/EMBLmyGFF3 |
| Flagger | v0.3.2 | https://github.com/mobinasri/flagger |
| winnowmap | 2.03 | https://github.com/marbl/Winnowmap |
| Secphase | v0.4.3 | https://github.com/mobinasri/secphase |
| DeepVariant | 1.4.0 | https://github.com/google/deepvariant |
| MUMmer | v4.0.0rc1 | https://github.com/mummer4/mummer |
| EMBOSS | 6.6.0 | https://emboss.sourceforge.net/ |
| OrthoFinder | 2.5.5 | https://github.com/davidemms/OrthoFinder |
| MAFFT | 7.526 | https://mafft.cbrc.jp/alignment/software/ |
| IQ-TREE | 2.3.6 | http://www.iqtree.org/ |
| ASTRAL-Pro3 | 1.16.2.4 | https://github.com/chaoszhang/ASTER |
| MCscanX | commit b1ca533 | https://github.com/wyp1125/MCScanX |
| Synvisio | commit 3415935 | https://synvisio.usask.ca/#/ |

**Supplementary Table 2: Different metrics based on alignment of pseudo-haplotype one of *L. fluviatilis* and *L. planeri* to each other and to a *L. fluviatilis* individual from the UK.**

| Aligned bases (percentage of genome assembly) | | | | | |
|---|---|---|---|---|---|
| | kcLamFluv2.2.h1 | kcLamFluv2.2.h2 | kcLamPlan1.2.h1 | kcLamPlan1.2.h2 | kcLamFluv1.1 |
| kcLamFluv2.2.h1 | | 881,074,207 (95.3675%) | 949,977,127 (90.5964%) | 870,262,887 (90.6182%) | 927,780,578 (88.9979%) |
| kcLamFluv2.2.h2 | 887,187,821 (91.3989%) | | 910,159,027 (86.7991%) | 854,622,156 (88.9896%) | 891,771,656 (85.5437%) |
| kcLamPlan1.2.h1 | 958,093,979 (89.2691%) | 901,211,315 (93.5558%) | | 888,105,065 (92.4760%) | 923,055,679 (88.5446%) |
| kcLamPlan1.2.h2 | 895,233,107 (83.4121%) | 863,916,684 (89.6842%) | 904,098,670 (86.2211%) | | 865,171,924 (82.9921%) |
| kcLamFluv1.1 | 952,584,705 (88.7558%) | 898,277,271 (93.2512%) | 940,046,391 (89.6493%) | 863,424,814 (89.9062%) | |
| **Insertions (sum in bp)** | | | | | |
| | kcLamFluv2.2.h1 | kcLamFluv2.2.h2 | kcLamPlan1.2.h1 | kcLamPlan1.2.h2 | kcLamFluv1.1 |
| kcLamFluv2.2.h1 | | 45,000 (96,896,895) | 71,559 (179,071,556) | 54,535 (117,194,461) | 68,502 (176,805,416) |
| kcLamFluv2.2.h2 | 48,389 (145,224,334) | | 66,059 (197,774,874) | 53,019 (128,383,196) | 62,507 (199,484,327) |
| kcLamPlan1.2.h1 | 74,939 (208,858,838) | 60,086 (134,891,062) | | 36,374 (83,167,882) | 68,711 (179,916,524) |
| kcLamPlan1.2.h2 | 67,161 (253,155,373) | 56,495 (169,199,520) | 45,401 (183,416,744) | | 62,646 (228,414,149) |
| kcLamFluv1.1 | 77,251 (216,215,970) | 61,269 (139,946,475) | 74,769 (190,651,971) | 57,777 (127,606,908) | |
| **SNPs** | | | | | |
| | kcLamFluv2.2.h1 | kcLamFluv2.2.h2 | kcLamPlan1.2.h1 | kcLamPlan1.2.h2 | kcLamFluv1.1 |
| kcLamFluv2.2.h1 | | 3,151,196 | 4,547,241 | 3,830,734 | 4,604,025 |
| kcLamFluv2.2.h2 | 3,151,196 | | 4,037,794 | 3,616,417 | 4,029,994 |
| kcLamPlan1.2.h1 | 4,547,241 | 4,037,794 | | 2,452,660 | 4,539,518 |
| kcLamPlan1.2.h2 | 3,830,734 | 3,616,417 | 2,452,660 | | 3,819,821 |

| | | | | | |
|---|---|---|---|---|---|
| kcLamFluv1.1 | 4,604,025 | 4,029,994 | 4,539,518 | 3,819,821 | |
| **Indels** | | | | | |
| | kcLamFluv2.2.h1 | kcLamFluv2.2.h2 | kcLamPlan1.2.h1 | kcLamPlan1.2.h2 | kcLamFluv1.1 |
| kcLamFluv2.2.h1 | | 4,021,416 | 5,879,800 | 4,833,123 | 5,949,776 |
| kcLamFluv2.2.h2 | 4,021,416 | | 5,144,900 | 4,517,162 | 5,129,985 |
| kcLamPlan1.2.h1 | 5,879,800 | 5,144,900 | | 3,138,986 | 5,885,164 |
| kcLamPlan1.2.h2 | 4,833,123 | 4,517,162 | 3,138,986 | | 4,837,701 |
| kcLamFluv1.1 | 5,949,776 | 5,129,985 | 5,885,164 | 4,837,701 | |
